# The *E. coli* CNF1 toxin induces fetal reprogramming of intestinal stem cells and a serrated colorectal cancer transcriptional signature in organoids

**DOI:** 10.1101/2025.10.02.680056

**Authors:** Diane Letourneur, Emma Bergsten, Nathan Raynaud, Juliette Mondy, Giulia Nigro, Landry-Laure Tsoumtsa Meda, Amaury Vaysse, Stéphane Descorps-Declere, Anna Yoshida, Caroline Stefani, Iradj Sobhani, Emmanuel Lemichez, Amel Mettouchi

## Abstract

Mechanistic insights are essential to understand how gut microbiota pathobionts contribute to tumorigenesis, given the mounting evidence linking them to colorectal cancer. We report a higher prevalence of the Rho GTPases-targeting *cnf1* toxin gene from *Escherichia coli* in the microbiota of early-stage, proximal colorectal cancer. Comparative gene set enrichment analysis reveals a concordant serrated pathway signature between colorectal cancer tissue of patients colonized with *cnf1*^+^ bacteria and CNF1-treated mouse intestinal organoids. RNA sequencing of organoids shows that CNF1 induces a fetal-like transcriptional reprogramming. Using integrated approaches, we demonstrate that CNF1 reprograms Lgr5⁺ stem cells into a Ly6a/Sca-1⁺ fetal-like state, that exhibits enhanced stemness potential. This reprogramming is preceded by a Yap/Taz-driven early transcriptional program and nuclear translocation of Yap. Functional analyses identify a RhoA/Rock–Yap/Taz–dependent transition to Ly6a/Sca-1⁺ stem cells, highlighting a mechanistic link between bacterial effectors and stem cell plasticity in colorectal tumorigenesis.

## INTRODUCTION

Colorectal cancer (CRC) remains a leading cause of cancer-related mortality worldwide. This multifactorial disease involves both intrinsic genetic cues and contribution of lifestyle and environmental factors. Over the past decade, microbiota composition and dysbiosis has been the center of interest in CRC research. Bacterial dysbiosis can promote chronic inflammation, a well-established driver of CRC initiation ^1^. Besides, bacterial dysbiosis allows emergence of specific pathobionts that directly participate in the process of carcinogenesis through diverse molecular mechanisms ^2,3^. CRC represents a heterogeneous malignancy and its pathogenesis depends on the anatomical location of the tumors, which show distinct histological and molecular features. The conventional model of adenoma-to-carcinoma sequence initiates with mutations in *APC* and stepwise acquisition of mutations in *KRAS* and *TP53* throughout life and accounts for the majority of CRCs. In an alternative pathway, a subset of CRCs arises from serrated polyps as precursor lesions that will eventually progress to cancer, the latter being more commonly observed in proximal (right-sided) colon ^4,5^.

Among pathobionts arising upon disruption of the gut microbial ecosystem, pathogenic strains of *Escherichia coli*, particularly adherent-invasive *E. coli* (AIEC) and extraintestinal pathogenic *E. coli* (ExPEC), have emerged as potential contributors to CRC development ^2,3,6,7^. A breakthrough in this field was the discovery of colibactin (Clb), a genotoxic metabolite synthesized by enzymes encoded within the polyketide synthase *pks* genomic island. Colibactin alkylates DNA and induces double-strand breaks, leading to a distinctive mutational signature. Such signature is found in healthy human colon crypts and with higher frequency in colorectal tumors, including within CRC driver genes, as assessed for the most commonly mutated gene *APC*, which contains up to 5.3% mutations matching the pks mutational signature ^8,9^, and *BRAF* ^10^. These seminal studies thereby implicate an *E. coli* toxin in the molecular etiology of CRC. Notably, up to 67% of CRC tissues are colonized by *E. coli* harboring the *pks* island compared to less than 35% of control biopsies ^11–14^. Colibactin-producing *pks* operon is highly prevalent in pathogenic strains of *E. coli* from the phylogenetic group B2, including ExPEC which possess enhanced capabilities for gut colonization, causing elevated risks of extra-intestinal infections. ExPEC strains have been isolated from the colonic mucosa of CRC patients ^15,16^. A significant proportion of ExPEC produce the Cytotoxic Necrotizing Factor 1 toxin (CNF1) together with colibactin ^12,13,17^. Recent findings reported that CNF1 endows *E. coli* with enhanced capacities for gut colonization, in conjunction with epithelial tissue invasion facilitated by its deamidase activity ^18^.

CNF1 is an AB-like toxin that binds to the ubiquitously expressed Lu/BCAM receptor at the host cells surface and translocates its catalytic domain into the cytosol via endosomal compartments ^19,20^. There, it catalyzes the deamidation of Rho GTPases—specifically glutamine 63 in RhoA (Q61 in Rac1 and Cdc42)—resulting in their constitutive activation, thereby disrupting important cellular functions. CNF1 provides an example of an environmental factor able to catalyze post-translational somatic modification of Rho GTPases ^21,22^. This deamidation blocks the intrinsic and GAP-stimulated GTPase activity, thereby leaving the Rho proteins permanently activated and able to transduce signaling cascades. Nevertheless, the activated forms of RhoA and Rac1 GTPases are secondarily targeted to ubiquitin-mediated proteasomal degradation by the Smurf1 ^23,24^ and Hace1 ^25,26^ E3 ubiquitin ligases respectively, thereby mitigating cytotoxic effects. Although Rac1 activation by CNF1 was shown to induce reactive oxygen species production by NADPH oxidase ^27^ and drive bacterial invasion into host tissues ^22,28,29^, little is known on the function of RhoA activation by CNF1 in host-pathogen interaction. Intriguingly, although data remain limited, scattered studies report that CNF1-producing *E. coli* are more frequently associated with mucosa of CRC patients compared to controls, with approximately 40% of CRC biopsies being colonized by CNF1-producing *E. coli versus* 12-13% of mucosa from controls ^11–13^. However, assessment of CNF1 carcinogenic potential in animal models gave divergent results with a pro-tumoral function in prostate cancer and colitis-associated CRC ^30,31^ or a protective role in the *Apc^Min/+^*mouse model of CRC ^32^. Therefore, further studies are warranted to clarify the context-dependent effects and underlying mechanisms.

In recent years, intestinal organoids have emerged as a powerful model system for investigating host–microbe interactions. Derived from adult stem cells, these three-dimensional epithelial structures faithfully mimic the cellular diversity, and spatial organization of the intestinal mucosa ^33^. This complexity renders them especially well-suited for studying how pathobionts interact with the host epithelium in a controlled yet biologically relevant context. The present study aimed to investigate CNF1-mediated alterations in the intestinal epithelium, using murine intestinal organoids. Here we demonstrate that exposure to the CNF1 toxin elicits a transcriptional program reminiscent of serrated CRC in primary intestinal organoids, accompanied by a reprograming of stem cells towards a fetal-like stem cell identity and increased stemness potential. Mechanistically, induction of fetal markers is driven by a RhoA-Yap/Taz signaling axis. In parallel, we report a significant enrichment of CNF1-expressing bacteria within the fecal microbiota of individuals harboring premalignant lesions and early-stage tumors in the proximal colon together with enrichment of serrated CRC signatures in RNAseq from *cnf1+* tumors, suggesting a potential role for CNF1-producing pathobionts in tumor initiation possibly through modulation of epithelial plasticity.

## RESULTS

### Increased prevalence of CNF1 and colibactin toxins-encoding genes in colorectal cancer-associated microbiota

In initiating this study, we analyzed seven publicly available shotgun metagenomic colorectal cancer datasets of human fecal microbiota to investigate the presence of the *cnf1*gene, encoding a Rho GTPase deamidase, and the *pks* genomic island spanning *clbA* to *clbS* genes (see Table 1 and Materials and Methods). These datasets originated from studies conducted across three continents—Europe, America, and Asia—and involved various sampling strategies and DNA extraction protocols ^34–40^. Our computational analysis encompassed 382 individuals with normal colonoscopy, 116 with adenomas, and 386 with colorectal cancer. Additionally, we performed qPCR analysis on material generated from stools and tissue-associated microbiota from cohorts located in France, designed FR2 and FR3, respectively, in Tables 1 and 2^34,41^. FR2 consists of 120 controls, 24 adenomas and 107 CRC samples and FR3 of 52 Tumor and Normal adjacent paired mucosal samples.

**Table 1.** List of the cohorts analyzed in this study.

| Cohort | Country | Project ID | Control | Adenoma | CRC | Analysis | Reference of the cohort |
| --- | --- | --- | --- | --- | --- | --- | --- |
| FR1 | France | PRJEB6070 | 61 | 42 | 53 | Computational | Zeller *et al.* 2014 <sup>35</sup>,<br>"study population F" |
| AT | Austria | PRJEB7774 | 63 | 47 | 46 | Computational | Feng *et al.* 2015 <sup>36</sup> |
| CN | China | PRJEB10878 | 54 | 0 | 74 | Computational | Yu *et al.* 2017 <sup>37</sup> |
| US | USA | PRJEB12449 | 52 | 0 | 52 | Computational | Vogtmann *et al.* 2016 <sup>38</sup> |
| DE | Germany | PRJEB27928 | 60 | 0 | 60 | Computational | Wirbel *et al.* 2019 <sup>40</sup> |
| IT | Italy | PRJNA447983 | 52 | 27 | 61 | Computational | Thomas *et al.* 2019 <sup>39</sup> |
| JP | Japan | PRJDB4176 | 40 | 0 | 40 | Computational | Wirbel *et al.* 2019 <sup>40</sup> |
| FR2 | France | — | 120 | 24 | 107 | qPCR stool | Sobhani *et al.* 2019<br>and this study <sup>34</sup> |
| FR3 | France | — | 0 | 0 | 52 | qPCR tissue | Sobhani *et al.* 2019<br>and this study <sup>34</sup> |

**Table 2.**
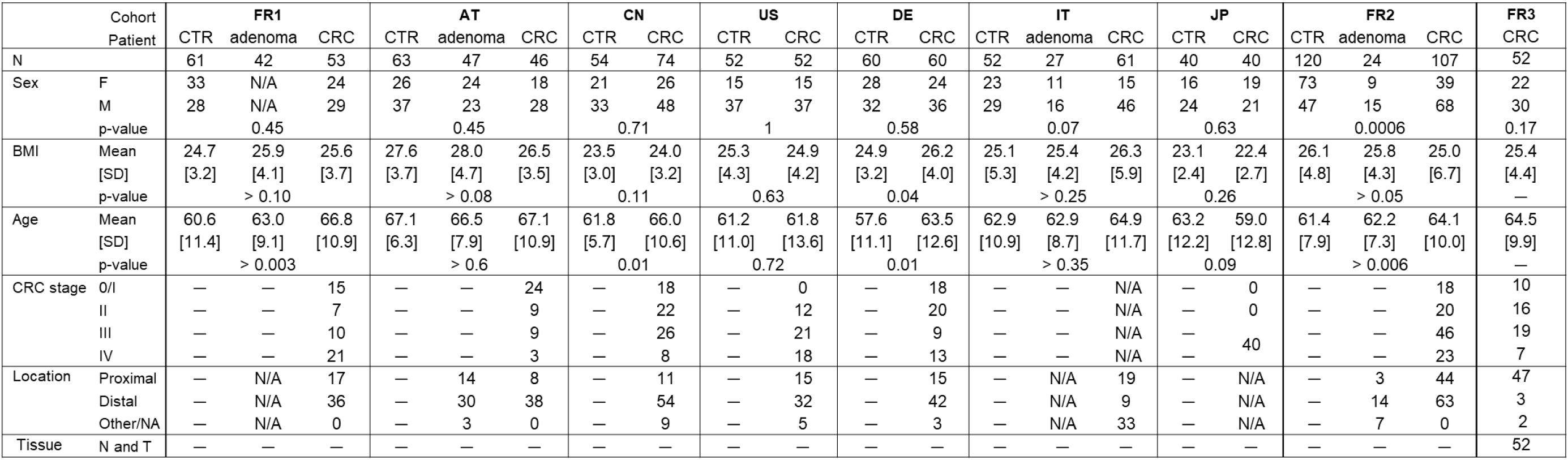
Table 2 summarizes the clinical and demographic characteristics of the cohorts analyzed in this study, including age, sex, body mass index (BMI), tumor stage and tumor location.

For the development of our computational pipeline (Figure S1A), we used the Diamond program, allowing the high throughput alignment of DNA reads against custom reference databases (see Materials and Methods section) ^42^. We detected *cnf1* in 16% of CRC (n=63/386) and 11% of adenomas (n=13/116), *versus* 9% (n=34/382) of controls, with a statistically significant 1.8 fold enrichment in CRC compared to controls (CRC *vs* controls: p < 0.01) (Figure 1A). *pks* was detected in 32% of CRC (n=123/386) and 26% of adenomas (n=30/116), *versus* 18% of controls (n=69/382), with a 1.8 fold statistically significant enrichment in CRC compared to controls (CRC *vs* controls: p < 0.0001) (Figure 1A). We then set out to detect *cnf1* and *clbB* (from the *pks* island) genes by qPCR in the FR2 cohort. In good agreement with the computational approach, we could detect a similar 1.7-fold increased prevalence of *cnf1* in CRC fecal microbiota as compared to controls, a trend detected at all stages of tumor development (Figure 1B, Table 3). *pks* detection by qPCR also mirrors the profile obtained by computational analysis, although the rate of detection is higher in all groups, with a 1.8-fold increase in prevalence within CRC microbiota as compared to the control group (Figure 1B, Table 3). Notably, when segregating FR2 patients according to CRC stage and location (proximal *vs* distal), we distinguished a higher prevalence of *cnf1* (Figure 1C, Table 3) and *pks* (Figure S1B, Table 3) in proximal stage I CRC, and similar trend for proximal adenoma, as compared to the prevalence in the control population. All patients tested positive for fecal *E. coli*, and *E. coli* levels did not significantly vary with CRC stage and location (Figure 1D). Higher detection rate of *cnf1* and *pks* in proximal stage I CRC is therefore not linked to higher *E. coli* colonization. Since the *pks* island is also present in *Klebsiella* species, we detected fecal *Klebsiella* by qPCR in the FR2 cohort and observed a tendency toward increased *Klebsiella* carriage in adenoma and CRC patients, with 22% (n=24/107) in CRC and 21% (n=5/24) in adenomas *versus* 13% (n=15/120) in controls (CRC *vs* controls: p = 0.07) and no increased prevalence in stage I proximal CRC (Table 3, Figure S1C). Interestingly, the bias towards early-stage proximal CRC association for *cnf1^+^ E. coli* was not echoed when the fecal carriage of two well-established cancer-associated bacteria, *Parvimonas micra* and *Fusobacterium nucleatum*, was assessed by Périchon and collaborators (personal communication) ^43^ (Figure S1D, S1E). We then examined if *cnf1*^+^ and *pks*^+^ bacteria colonize CRC colonic mucosa by qPCR detection of their genes in DNA extracted from tissue of paired CRC specimen of the FR3 cohort. *cnf1* (Figure 1E) and *pks* (Figure 1F) were detected in more than 25% of tumoral (T) and adjacent non-tumoral tissues (N) with no significant difference between T and N. This indicates that bacteria encoding these toxins are found in association with the colonic tissue, an important feature for potential carcinogenic impacts on the host.

**Figure 1.**
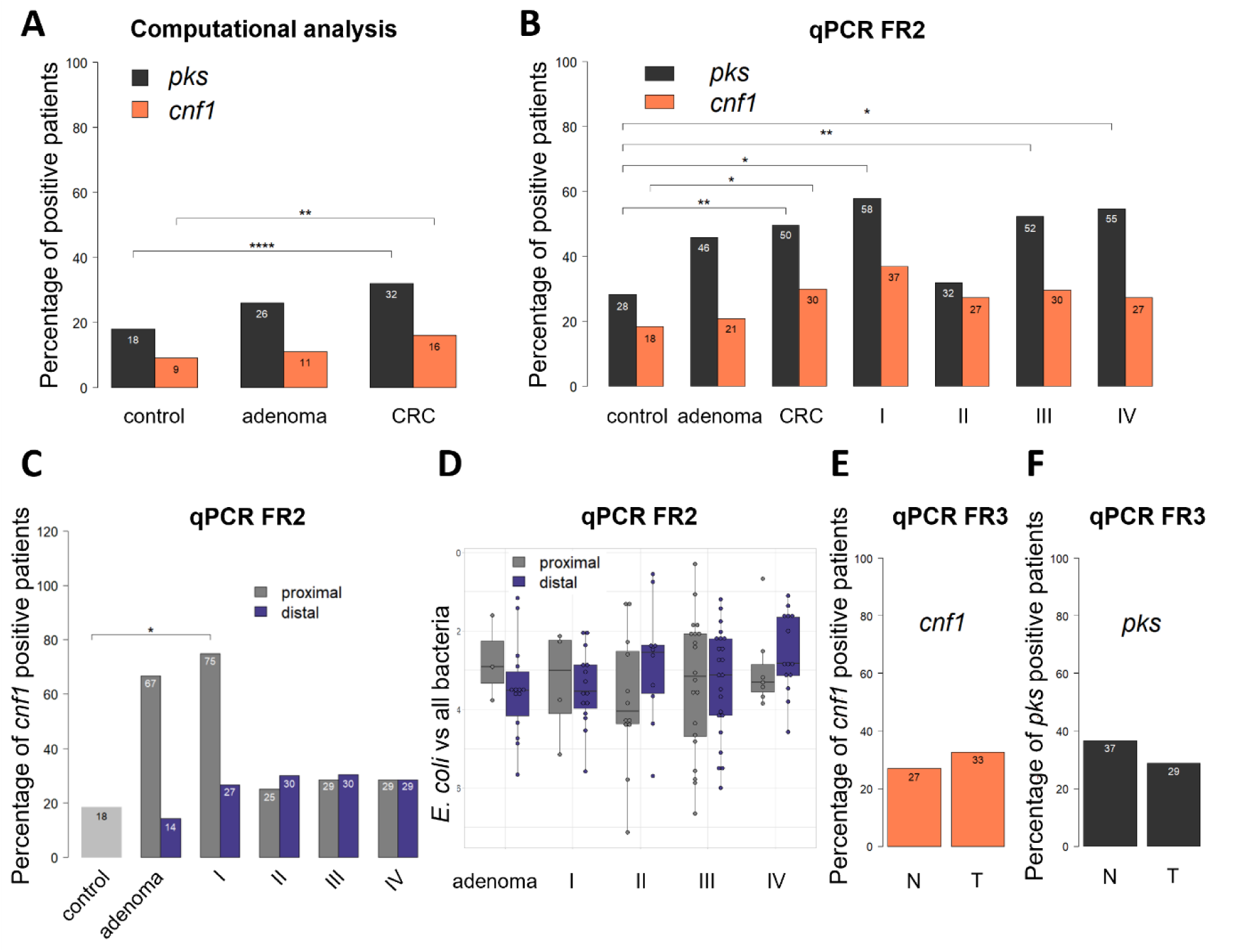
Increased prevalence of CNF1 and Colibactin toxins-encoding genes in colorectal cancer-associated microbiota. (A-B) Percentage of patients with fecal carriage of *cnf1* and *pks,* determined by bioinformatics on WGS of the FR1, AT, CN, US, DE, IT, and JP cohorts (A) and by qPCR in the FR2 cohort (B). Chi2 and Fisher’s exact test were used to determine statistical significance pairwise. ^∗∗∗∗^ *p* < 0.0001, ^∗∗^ *p* < 0.01, ^∗^ *p* < 0.05. (C) Percentage of FR2 patients with fecal carriage of *cnf1*, depending on the stage and tumor location. Chi2 or Fisher’s exact test were used to determine statistical significance by pairs. ^∗^ *p* < 0.05. (D) Fecal abundance of *E. coli* relatively to all bacteria measured by qPCR in patients of the FR2 cohort. Data are represented as median with interquartile range, with each dot representing one patient. Statistical significance was assessed using pairwise Wilcoxon rank-sum test with Benjamini-Hochberg correction; not significant. (E-F) Percentage of carriage of *cnf1* (E) and *pks* (F) in tumoral (T) and adjacent non tumoral (N) tissues from FR3 patients. Fisher’s exact test was used to determine statistical significance; not significant.

**Table 3.** Table 3 summarizes the number of patients from the FR2 cohort harboring *cnf1* detection (*cnf1*^+^), ClbB detection (*pks*^+^), and *Klebsiella* spp. detection within the fecal microbiota (*Klebsiella*^+^). Data are represented by groups and in function of tumor location or stage. The information for location was missing for one patient in the CRC stage-IV group and for 7 patients in the adenoma group.

|  |  | *pks+* | *cnf1+* | *Klebsiella+* | n total |
| --- | --- | --- | --- | --- | --- |
| Control | total | 34 (28%) | 22 (18%) | 15 (12.5%) | 120 |
| adenoma | proximal | 2 (67%) | 2 (67%) | 1 (33.3%) | 3 |
|  | distal | 7 (50%) | 2 (14%) | 3 (21.4%) | 14 |
|  | total | 11 (46%) | 5 (21%) | 5 (20.8%) | 24 |
| CRC | proximal | 17 (39%) | 14 (32%) | 9 (20.5%) | 44 |
|  | distal | 36 (58%) | 18 (29%) | 15 (24.2%) | 62 |
|  | total | 53 (50%) | 32 (30%) | 24 (22.4%) | 107 |
| I | proximal | 4 (100%) | 3 (75%) | 0 (0%) | 4 |
|  | distal | 7 (47%) | 4 (27%) | 4 (26.7%) | 15 |
|  | total | 11 (58%) | 7 (37%) | 4 (21.1%) | 19 |
| II | proximal | 2 (17%) | 3 (25%) | 1 (8.3%) | 12 |
|  | distal | 5 (50%) | 3 (30%) | 3 (30.0%) | 10 |
|  | total | 7 (32%) | 6 (27%) | 4 (18.2%) | 22 |
| III | proximal | 8 (38%) | 6 (29%) | 7 (33.3%) | 21 |
|  | distal | 15 (65%) | 7 (30%) | 7 (30.4%) | 23 |
|  | total | 23 (52%) | 13 (30%) | 14 (31.8%) | 44 |
| IV | proximal | 3 (43%) | 2 (29%) | 1 (14.3%) | 7 |
|  | distal | 9 (64%) | 4 (29%) | 1 (7.1%) | 14 |
|  | total | 12 (55%) | 6 (27%) | 2 (9.1%) | 22 |

Together, our integrative qPCR and metagenomic analyses across multiple international cohorts reveal a significant enrichment of *cnf1*-encoding and *pks*-harboring bacteria in the fecal microbiota of CRC patients, with a striking association of *cnf1* with proximal location at early tumor stage.

### The CNF1 toxin of *E. coli* triggers a transcriptional signature of serrated carcinogenesis in intestinal organoids

Intestinal organoids have emerged as a powerful *ex vivo* model system to study host–pathogen interactions and colorectal cancer development, offering a physiologically relevant model of cellular diversity of the intestinal epithelium and maintenance of a high turnover rate driven by stem cells of the crypt base (CBC) ^33^. To gain mechanistic insights on the link between *cnf1^+^*bacteria and CRC pathology, we studied the effect of the CNF1 toxin on murine intestinal organoids, using a combination of imaging and transcriptomics approaches. We recorded a high proliferative index as assessed by Ki67 immunolabeling and no significant changes in response to CNF1 treatment (Figure 2A, B). Interestingly, in organoids stimulated by CNF1, we observed lower expression of differentiation markers for enterocytes (*Anpep*, Figure S2A), Paneth cells (*Defa5*, Figure S2B), enteroendocrine cells (*Chga*, Figure S2C), and Tuft cells (*Dclk1*, Figure S2D), but not for goblet cells (*Muc2*, Figure S2E). We verified that the catalytically inactive mutant C866S of CNF1 toxin had no effect on the expression of these cell type-specific markers. This prompted us to characterize the molecular responses to CNF1, performing transcriptional analysis by RNA-seq at different times along a 48h period. We observed global changes at the transcriptional level with a number of differentially expressed genes (DEG) of 221 at 4h, 3590 at 24h and 8317 at 48h treatment (Figure 2C, Supplementary Information 1). We observed a dynamic response overtime and stepwise establishment of a genetic response, with only a minority (124) of DEG common to the three conditions and 3020 DEGs shared between 24h and 48h treatment. KEGG pathway enrichment analysis revealed that up-regulated genes at 24h were highly associated with pathways related to endocytosis and cornified envelope formation, that may reflect that intestinal cells harbor aberrant differentiation program in response to CNF1. At later timepoint, pathways involved in drug metabolism, chemical carcinogenesis, PPAR signaling, lipid metabolism and absorption, and host responses to infectious processes became enriched. These temporal dynamics are illustrated in Figure S2F which highlights the top 50 enriched KEGG pathways.

**Figure 2.**
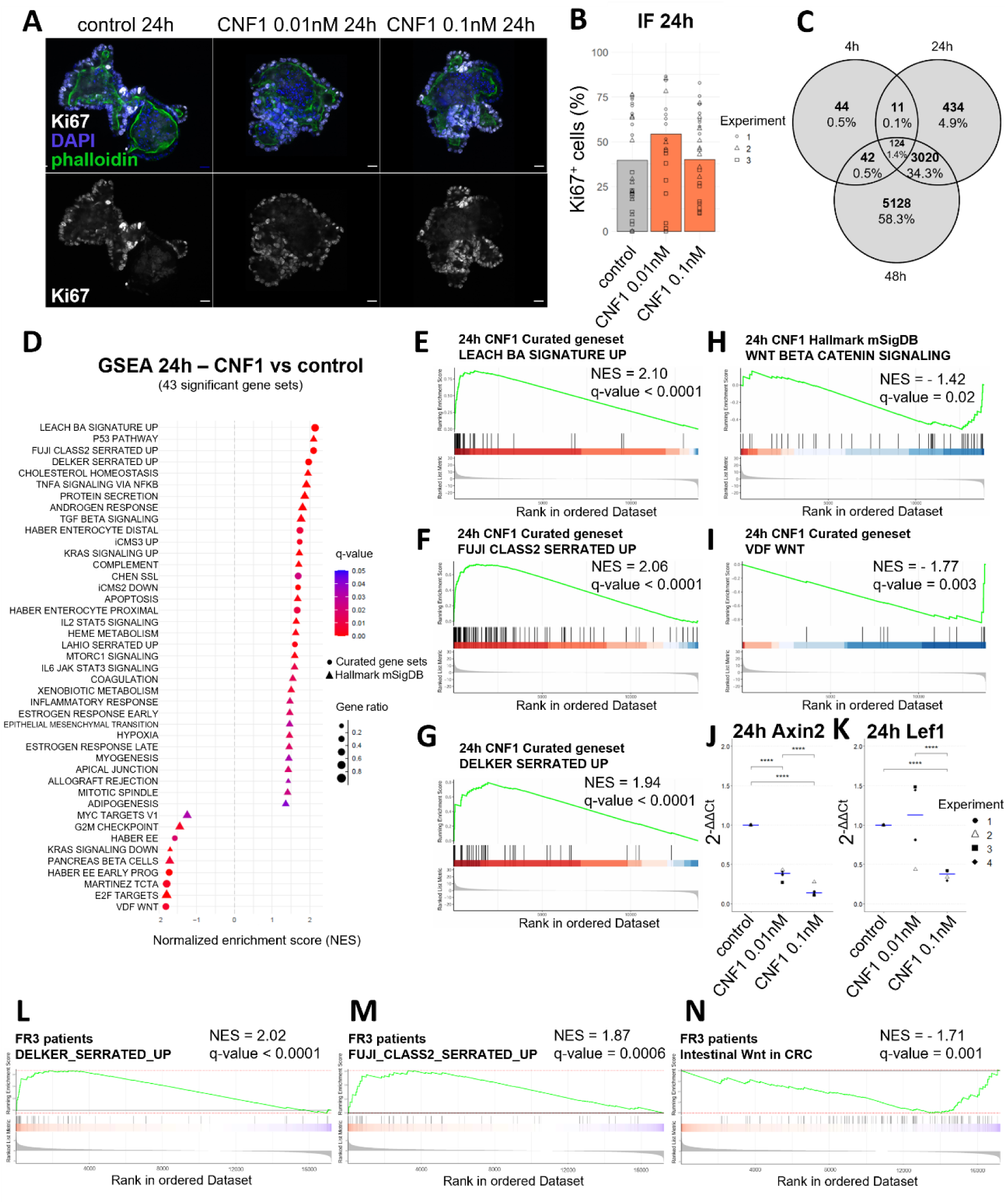
The CNF1 toxin of *E. coli* triggers a serrated pathway transcriptional signature of carcinogenesis in intestinal organoids. (A) Immunofluorescence staining for proliferation in small intestinal organoids after 24h of CNF1 stimulation. Nuclei are stained with DAPI, actin with phalloidin, and proliferative cells with Ki67. Scale bar: 20 µm. (B) Percentage of Ki67-positive cells determined by immunofluorescence after 24h of CNF1 stimulation. Data are represented as a mean (3-11 organoids/condition/experiment from n= 3 independent experiments). Statistical significance was determined by generalized linear regression using independent experiments as random effects with Tukey’s correction; not significant. (C) Venn diagram of differentially expressed genes in small intestinal organoids treated for 4h, 24h, or 48h with 0.1nM of CNF1 (n=3 independent experiments). Differentially expressed genes number (in bold) and percentage over the total 8803 differentially expressed genes overtime are indicated. Differential analysis and statistical tests were performed with DESeq, with Benjamini-Hochberg correction. (D) GSEA analysis ranking published signatures (“curated genesets”) and mSigDB hallmark gene sets according to normalized enrichment score (NES) and q-value by Monte Carlo analysis with Benjamini-Hochberg correction (n=3 independent experiments). Organoids treated 24h with 0.1 nM of CNF1 are compared to non-treated controls. (E-I) GSEA analysis in organoids after 24h of 0.1 nM CNF1 stimulation showing the Delker signature of sessile serrated adenomas and polyps (E), the Fuji signature of human organoids from serrated lesions (F), the Leach signature of a mouse model of proximal CRC (G), the mSigDB hallmark of Wnt/β-catenin signaling (H), and the Van der Flier Wnt signaling signature in CRC (I). Statistical significance was determined by Monte Carlo analysis with Benjamini-Hochberg correction (n=3 independent experiments). (J-K) RTqPCR showing expression of the Wnt effectors *Axin2* (H) and *Lef1* (I) after 24h of CNF1 stimulation, with median in blue (n=4 independent experiments). Statistical significance was determined by generalized linear regression using independent experiments as random effects, with Tukey’s correction. ^∗∗∗∗^ *p* < 0.0001. (L-N) GSEA analysis of patients from the FR3 cohort with *cnf1* tissue carriage compared to patients without *cnf1* carriage, showing the Delker signature of sessile serrated adenomas and polyps (L), the Fuji signature of human organoids from serrated lesions (M) and the Van der Flier Wnt signaling signature in adenoma and CRC (N). Statistical significance was determined by Monte Carlo analysis with Benjamini-Hochberg correction.

To gain complementary functional insights into the transcriptional changes observed in our RNA-seq data, we performed Gene Set Enrichment Analysis (GSEA), which enables the identification of biologically relevant pathways and processes associated with coordinated gene expression changes. For this, we screened the 50 hallmark gene sets within mouse molecular signature databases (mSigDB) collections together with 35 implemented curated gene sets previously published representing signatures of the conventional adenoma-to-carcinoma pathway of colon tumorigenesis and signatures of the alternative serrated pathway of colorectal cancer, as well as signatures of differentiated epithelial cells (listed in Supplementary Information 2). Figure 2D shows the ranking of signatures present in the gene expression profile of organoids after 24h of CNF1 treatment. Interestingly, the transcriptional response shows marked enrichment of serrated pathway-related signatures, that rank in the top 4 gene sets of the list. Indeed, the Leach signature ^44^ defined in a mouse model of proximal-CRC associated with *Braf^V^*^600^*^E^* activating mutation and loss of epithelial TGF-beta receptor signaling displays a robust and significant positive enrichment (NES = 2.1, q < 0.0001; Figure 2E) in CNF1-treated organoids. Similar enrichment is observed with the Fuji signature ^45^, defined from organoids cultured from human serrated lesions (NES = 2.06, q < 0.0001; Figure 2F) or the Delker signature ^45^, generated from human sessile serrated adenomas and polyps (NES = 1.94, q < 0.0001; Figure 2G). Conversely, signatures of conventional CRC ^46,47^ are not enriched in CNF1-treated organoids (Figure S2G, H).

Consistent with an unrestrained Wnt/β-catenin pathway being the main initiator event of conventional CRC, we obtain negative enrichment score for the hallmark-Wnt/β-catenin signaling gene set or the Van der Flier gene set for intestinal Wnt signaling (NES = −1.42, q = 0.02 and NES = −1.77, q = 0.003 respectively; Figure 2H, I). We could confirm the default of activation of the Wnt pathway in CNF1-treated organoids by RT-qPCR measure of expression of the β-catenin target genes *Axin2* and *Lef1* (Figure 2J, K).

We leveraged RNA-seq data generated from patient tissue of the FR3 cohort by Bergsten *et al.* ^41^ to bridge mechanistic RNA-seq data from mouse organoids with epidemiological RNA-seq profiles from human tissues. We assessed enrichment of serrated neoplasia or Wnt transcriptional signatures by GSEA, comparing CRC patients harboring *cnf1*-encoding bacteria mucosal colonization with *cnf1*-negative CRC patients. Interestingly, mucosa from *cnf1*-positive CRC patients display a robust and significant positive enrichment in the Delker signature ^45^ for sessile serrated adenomas and polyps (NES = 2.02, q < 0.0001; Figure 2L) and the Fuji signature ^45^ of organoids cultured from human serrated lesions (NES = 1.87, q = 0.0006; Figure 2M). Conversely, the Van der Flier Wnt signaling signature in adenoma and CRC ^48^ showed a significantly negative enrichment, suggesting preferential downregulation in mucosa from *cnf1*-positive CRC patients (NES = −1.71, q < 0.001; Figure 2N). The Chen signature of conventional colon adenocarcinoma ^47^ reached no significant enrichment (NES = −0.90, q = 0.75; Figure S2I). These findings are in strong agreement with our observations in CNF1-treated organoids, thus reinforcing the relevance of the mechanistic insights gained from our combined approaches.

Altogether, we uncover that CNF1 induces changes in the intestinal epithelium leading to enrichment of signatures associated with serrated tumorigenesis, suggesting involvement of this alternative pathway in carcinogenesis associated to gut carriage of *cnf1^+^* bacteria.

### The CNF1 toxin of *E. coli* activates a fetal-like transcriptional program in stem cells

The serrated pathway of colorectal tumorigenesis has been previously associated with apparent lack or attenuation of Wnt pathway activation combined with high expression of fetal markers ^44,45^. We therefore implemented our DEG analysis in response to CNF1 by searching for enrichment in expression of genes of the Muñoz signature for Lgr5^+^ intestinal stem cells ^49^, the Mustata signature of fetal intestinal self-renewing spheroids ^46^ and the Pikkupeura signature of fetal proximal murine small intestine ^50^. The volcano plot in Figure 3A highlights within transcriptional response to CNF1 the DEGs reported in the Muñoz (in blue) and Mustata (in red) genesets. Remarkably, we observed a segregation of these signatures in two opposite groups, with a downregulation of markers of the Lgr5^+^ population, including *Lgr5* itself and *Olfm4* genes, and an upregulation of Lgr5-negative fetal stem cells markers including the well-established *Ly6a* (encoding Sca-1) and *Clu* genes. GSEA analysis using these gene sets confirms enrichment in fetal stem cell transcriptomic program (NES = 2.58, q < 0.0001 and NES = 2.00, q < 0.0001 for Mustata and Pikkupeura signatures respectively; Figure S3A, B) within the transcriptome of CNF1-stimulated organoids and enrichment in adult intestinal stem cell gene program in control organoids (NES = −2.97, q < 0.0001; Figure S3C).

**Figure 3.**
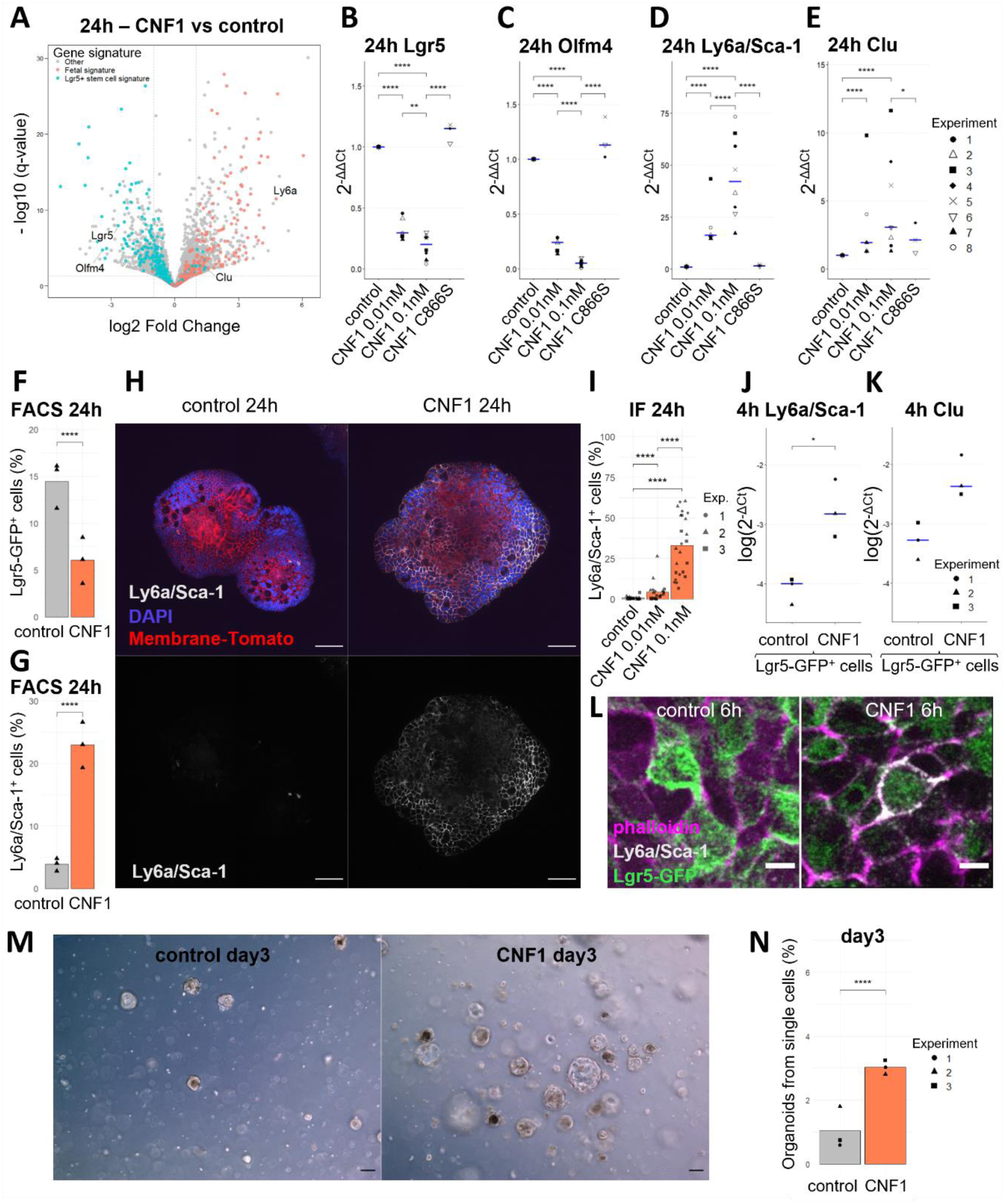
The CNF1 toxin of *E. coli* activates a fetal-like transcriptional program in stem cells. (A) Volcano plot showing differential expression analysis of small intestinal organoids after 24h of stimulation with 0.1 nM of CNF1. Markers of crypt based columnar cells (CBC) from the Muñoz signature of Lgr5^+^ cells are represented in red, and markers of the fetal intestinal epithelium from the Mustata signature are in blue. Differential analysis and statistical tests were performed with DESeq, with Benjamini-Hochberg correction (n=3 independent experiments). (B-E) RTqPCR showing expression of the Lgr5^+^ stem cell markers *Lgr5* (B) and *Olfm4* (C), and the fetal markers *Ly6a*/Sca-1 (D) and *Clu* (E) in small intestinal organoids after 24h of stimulation with WT or C866S mutant CNF1 (catalytically inactive), with median in blue (n=8 independent experiments). Statistical significance was determined by generalized linear regression using independent experiments as random effects, with Tukey’s correction. ^∗∗∗∗^ p < 0.0001, ^∗∗^ p < 0.01, ^∗^ p < 0.05. (F-G) Percentage of Lgr5-GFP^+^ (F) and Ly6a/Sca-1^+^ (G) cells sorted by flow cytometry. Data are represented as a mean (n=3 independent experiments). Statistical significance was determined by generalized linear regression using independent experiments as random effects. ^∗∗∗∗^ p < 0.0001. (H) Immunofluorescence staining of Ly6a/Sca-1 in organoids after 24h of 0.1 nM CNF1 stimulation. Nuclei are stained with DAPI, organoids from mT/mG mice express a membrane-targeted Tomato. Scale bar: 50 µm. (I) Quantification of the percentage of Ly6a/Sca-1^+^ cells determined by immunofluorescence. Data are represented as mean, with each dot representing an organoid (6-9 organoids/condition/experiment from n= 3 independent experiments). Statistical significance was determined by generalized linear regression using independent experiments as random effects, with Tukey’s correction. ^∗∗∗∗^ p < 0.0001. (J-K) RTqPCR showing expression of the fetal markers *Ly6a*/Sca-1(L) and *Clu* (M) in Lgr5-GFP^+^ cells sorted by FACS after 4h of 0.1 nM CNF1 stimulation, with median in blue (n=3 independent experiments). Statistical significance was determined by generalized linear regression using independent experiments as random effects. ^∗^ p < 0.05. (L) Immunofluorescence staining of Ly6a/Sca-1 in Lgr5-GFP organoids after 6h of 0.1 nM CNF1 stimulation. Actin is stained with phalloidin. Scale bar: 5 µm. (M) Organoid-forming capacity of cells dissociated from control organoids or organoids treated for 24h with 0.1 nM CNF1, and cultured for 3 days in control or CNF1-containing medium, respectively. Scale bar: 100 µm. (N) Quantification of organoid-forming capacity at day 3. Data are represented as a mean (n=3 independent experiments). Statistical significance was determined by generalized linear regression using independent experiments as random effects. ^∗∗∗∗^ p < 0.0001.

To confirm RNA-seq data, we performed RT-qPCR to quantify the expression of selected markers. CNF1 induced a dose-dependent decrease of *Lgr5* and *Olfm4* expression in small intestinal organoids (Figure 3B, C) as well as *Lgr5* and *Vdr* in colonic organoids - since *Olfm4* is not reliably expressed in the murine colon - (Figure S3D, E), dependent on its catalytic activity (see CNF1 C866S). In parallel, we observed a dose-dependent increase of *Ly6a*/Sca-1 and *Clu* expression in small intestine-derived organoids (Figure 3D, E) and in colonic organoids exposed to CNF1 (Figure S3F, G). These results were consolidated by measures on other fetal-like markers *Anxa1*, *Ly6d*, and *Krt7,* that showed coherent regulations (data not shown).

We validated these data at the cell population level by flow cytometry (FACS) analysis, using small intestinal organoids derived from *Lgr5-EGFP-IRES-CRE^ERT^*^2^ mice (Lgr5-GFP mice). FACS analysis (Figure S3H-J) revealed a two-fold decrease in the percentage of Lgr5-GFP^+^ cells (Figure 3F and Figure S3I) and a 6-fold increase in the proportion of Ly6a/Sca-1^+^ cells (Figure 3G and Figure S3J) after 24h of CNF1 treatment. CNF1-mediated expansion of Ly6a/Sca-1^+^ cells was confirmed by confocal microscopy (Figure 3H, I).

We then explored whether we could catch a window where Lgr5 identity is still present, and the fetal-like program initiates. Upon 4h treatment of organoids, we recorded increased levels of *Ly6a*/Sca-1 and *Clu* markers expression in the sorted Lgr5-GFP^+^ population (Figure 3J, K) indicating that Lgr5-GFP^+^ cells can directly give rise to the Ly6a/Sca-1^+^ population in response to CNF1-mediated cues. We controlled in parallel that Lgr5-GFP^+^ cells are not disappearing from the organoids upon cell death, as indicated by similar viability rate of treated and untreated GFP^+^ and GFP^-^ cells at this timing (Figure S3K). Confocal analyses corroborate the conclusion at the protein level, since we detect that Lgr5-GFP^+^ cells express Ly6a/Sca-1 at their surface upon 6h CNF1 treatment, confirming that Lgr5-GFP^+^ stem cells give rise to fetal-like Ly6a/Sca-1^+^ progenitors (Figure 3L). RNAscope targeting Lgr5 mRNA coupled to immunostaining of Ly6a/Sca-1 confirms in a WT background that Lgr5^+^ cells start expressing the Ly6a/Sca-1 marker at 6h while they gradually decrease their Lgr5 mRNA expression (Figure S3L, M) in response to CNF1.

To explore whether such activation of a fetal-like stem cell program leads to a fitness advantage, we evaluated the stemness potential of CNF1-treated organoids through monitoring of their organoid-forming ability. After 24h of CNF1 stimulation, cells dissociated from organoids were seeded in the presence of CNF1 for a further 3 days period, and organoid/spheroid formation was assessed in comparison of control non-treated organoids that were dissociated and seeded in parallel. More organoids were obtained from CNF1-treated conditions (Figure 3M, N), showing that the CNF1 toxin confers higher stemness potential.

Together, these experimental approaches converge to show that exposure to the Rho deamidase toxin CNF1 drives a fate change in Lgr5⁺ stem cells, leading to the acquisition of fetal markers, associated with enhanced stemness potential.

### Yap/Taz activity is enhanced by CNF1 in intestinal organoids, through a RhoA/Rock-dependent mechanism

Recent literature has implicated the Hippo-pathway effectors Yes-associated protein (Yap) and transcriptional co-activator with PDZ-binding motif (Taz) in the reprograming of epithelial cell populations towards a fetal-like state during irradiation and colitis-associated tissue damage and regeneration ^51–53^.

Indeed, at early time after CNF1 treatment, we identified in RNA-seq data a strong transcriptional activation of Yap/Taz effectors (Figure 4A) and GSEA enrichment of the Yap signature from Gregorieff ^52^, detectable along the time course (NES = 1.04, q = 0.0002 and NES = 1.44, q < 0.0001 at 4h and 24h respectively; Figure 4B, C). The Yap gene signature displayed targets either transiently induced or rising gradually at 24h and/or 48h (Figure S4A). Increased expression of the early Yap effectors *Ccn1*/*Cyr61* and *Ccn2*/*Ctgf* was verified by RT-qPCR in organoids derived from small intestine (Figure S4B-C) and colon (Figure S4D-E). In addition, Yap transcription factor activation state could be directly monitored by immunolabeling (Figure 4D-F), consistently showing 1.25-fold enhanced nuclear translocation upon 4h stimulation of organoids with CNF1 (Figure 4F). To formally establish the involvement of Yap/Taz in CNF1-triggered induction of a fetal-like stem cell transcriptional program, we used a chemical perturbation approach. Treatment of organoids with the Yap/Taz inhibitor verteporfin in the presence of CNF1 attenuated the induction of *Ly6a*/Sca-1 and *Clu* expression (Figure 4G, H), demonstrating contribution of Yap/Taz in our phenotype.

**Figure 4.**
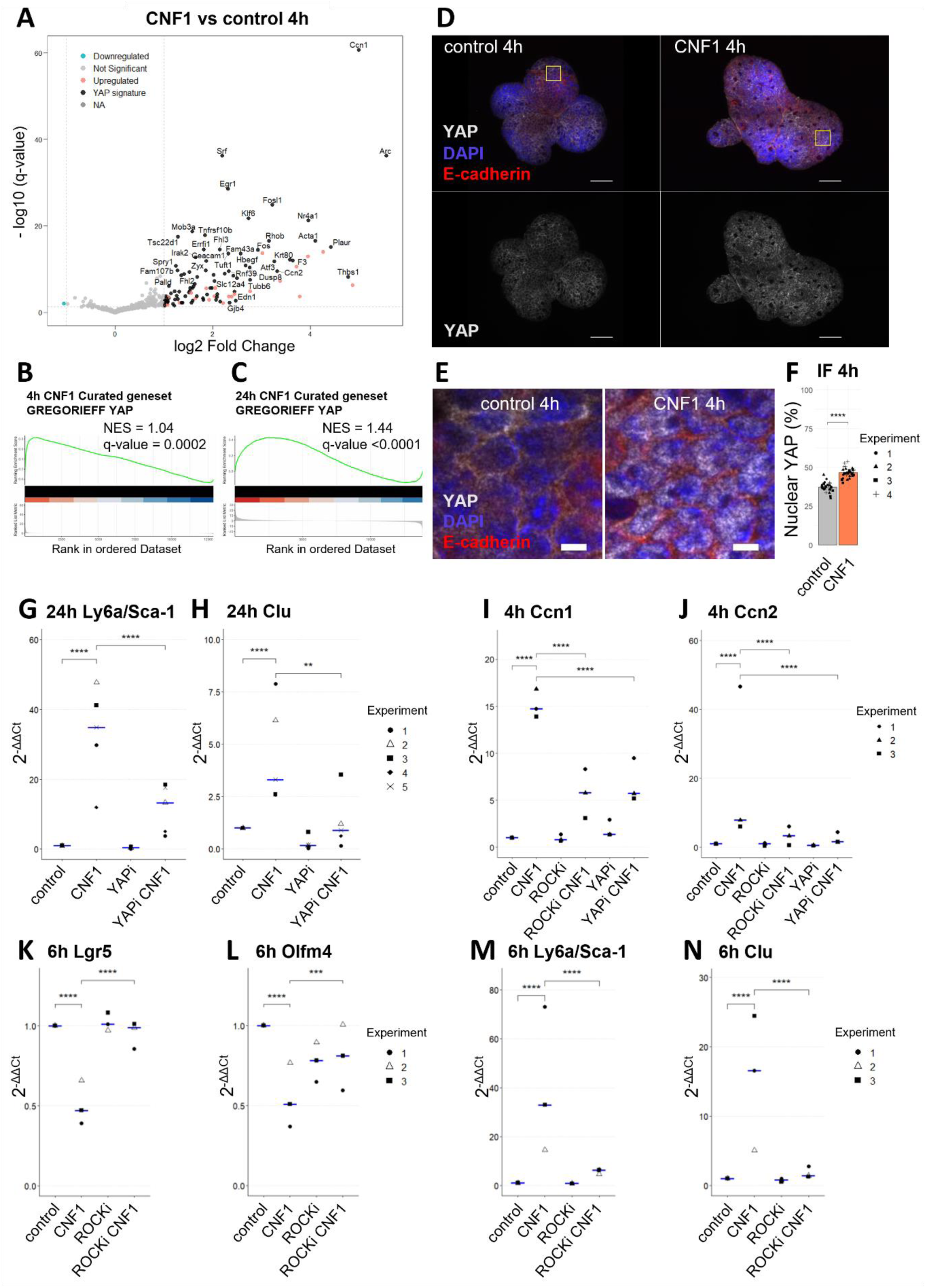
Yap/Taz activity is enhanced by CNF1 in intestinal organoids, in a RhoA/Rock-dependent mechanism. (A) Volcano plot showing differential expression analysis in organoids after 4h of stimulation with 0.1 nM of CNF1 (n=3 independent experiments). Previously described effectors of the Yap pathway and genes from the Gregorieff signature are represented in black. Differential analysis and statistical tests were performed with DESeq, with Benjamini-Hochberg correction. (B-C) GSEA analysis of the Gregorieff Yap signature after 4h (B) or 24h (C) of 0.1 nM CNF1 stimulation. Statistical significance was determined by Monte Carlo analysis with Benjamini-Hochberg correction (n=3 independent experiments). (D-E) Immunofluorescence staining of Yap nuclear translocation in small intestinal organoids after 4h of stimulation with 0.1 nM CNF1. Nuclei are stained with DAPI and E-Cadherin marks cell-cell junctions. Scale bar: 50 µm (D) or 5 µm (E). (F) Quantification of the percentage of cells with nuclear Yap determined by immunofluorescence (6-8 organoids/condition/experiment from n= 4 independent experiments). Statistical significance was determined by generalized linear regression using independent experiments as random effects. ^∗∗∗∗^ p < 0.0001. (G, H) RTqPCR showing expression of the fetal markers *Ly6a*/Sca-1 (G) and *Clu* (H) in organoids co-stimulated for 24h with 0.1 nM of CNF1 and a pharmacological inhibitor of Yap-Tead interaction (YAPi, verteporfin), with median in blue (n=5 independent experiments). Statistical significance was determined by generalized linear regression using independent experiments as random effects, with Tukey’s correction. ^∗∗∗∗^ p < 0.0001, ^∗∗^ p < 0.01. (I, J) RTqPCR showing expression of the Yap target genes *Ccn1*/*Cyr61* (I) and *Ccn2*/*Ctgf* (J) in organoids co-stimulated for 4h with 0.1 nM CNF1 and a pharmacological inhibitor of Yap-Tead interaction (YAPi, verteporfin) or Rock (ROCKi, Y-27632), with median in blue (n=3 independent experiments). Statistical significance was determined by generalized linear regression using independent experiments as random effects, with Tukey’s correction. ^∗∗∗∗^ p < 0.0001. (K-N) RTqPCR showing expression of *Lgr5* (K), *Olfm4* (L), *Ly6a*/Sca-1 (M), and *Clu* (N) in organoids co-stimulated for 6h with 0.1 nM of CNF1 and a pharmacological inhibitor of Rock (ROCKi, Y-27632), with median in blue (n=3 independent experiments). Statistical significance was determined by generalized linear regression using independent experiments as random effects, with Tukey’s correction. ^∗∗∗∗^ p < 0.0001, ^∗∗∗^ p < 0.001.

Because the CNF1 deamidase activates the RhoA GTPase ^21,22^, we investigated the involvement of the RhoA/Rock pathway in Yap/Taz activation downstream CNF1. Blocking the RhoA pathway with the Rock inhibitor Y-27632 impaired expression of *Ccn1*/*Cyr61* and *Ccn2*/*Ctgf* in response to CNF1 (Figure 4I, J), to the same extent as verteporfin, pointing towards RhoA-Rock involvement in Yap/Taz activation by CNF1. In agreement, the Rock inhibitor could abrogate CNF1-mediated decrease of *Lgr5* and *Olfm4* expression (Figure 4K, L) and increase of *Ly6a*/Sca-1 and *Clu* (Figure 4M, N) expression. Altogether, these findings indicate that CNF1-driven alterations in the fate of stem cell populations depend on the RhoA/Rock axis and Yap/Taz transcriptional co-activators.

## DISCUSSION

In this study, we set out to characterize the cellular effects of CNF1 on the gut epithelium. Using murine intestinal organoids, we show that CNF1 induces a serrated CRC transcriptional signature and reprograms epithelial stem cells toward a fetal-like identity, characterized by a RhoA–Yap/Taz-dependent induction of fetal markers. In parallel, we show that CNF1-expressing bacteria are enriched in the microbiota of patients with proximal premalignant lesions and proximal early-stage CRC.

CNF1, often produced by the same *E. coli* strains as the colibactin genotoxin, has been proposed as a pro-tumorigenic virulence factor, as enrichment of *cnf1*^+^ bacteria is detected in fecal and mucosal samples of CRC patients ^11–13,16,54^. Here, we detect increased prevalence of the CNF1 toxin-encoding gene within public metagenomic data across several cohorts investigating CRC-associated microbiota worldwide, encompassing 382 controls, 116 adenoma and 386 CRC. We could also confirm by our independent computational approach the enrichment of genes forming the *pks* island, responsible for colibactin synthesis within these cohorts, as previously assessed in ^55^. By generating refined *cnf1* and *pks* island gene databases that were used in a high throughput Diamond blast approach, we obtained enhanced detection rates for *cnf1*, that were not previously reached since a prior search in the same data base failed in its identification ^35^. Recently, a similar rationale allowed detection of several virulence factor genes (VFG) association with pathologic conditions, upon implementing the VFDB database with a broad spectrum of orthologous VFG sequences. Interestingly, the *E. coli* toxins CNF1 and alpha-hemolysin were the only disease-specific enriched VFG, restricted to CRC ^56^.

We could verify *cnf1* prevalence in CRC by quantitative PCR on fecal DNA from a French cohort. Moreover, we show in this cohort a clinical association of *cnf1*^+^ bacteria within the fecal microbiota of patients harboring pre-cancerous lesions and early-stage adenocarcinoma of the proximal colon, pointing towards a role of CNF1-producing *E. coli* in initiating events involved in proximal CRC. Previous studies reported no significant difference of *pks* fecal carriage depending on tumor location when considering all stages ^57,58^, and to our knowledge, fecal association of *cnf1*^+^ bacteria with proximal or distal CRC has not yet been explored. While colibactin can generate double strand breaks leading to mutations detected in tumors ^8,9,59^, evidence of CNF1 pro-carcinogenic function remain sparse. Our findings now extend this view by showing that CNF1 is not only associated epidemiologically with proximal CRC but also capable of reprogramming the intestinal epithelium in ways consistent with serrated tumorigenesis. Of note, the serrated tumorigenesis pathway is often associated with proximal CRC ^60^, which is consistent with our epidemiological study showing *cnf1* prevalence in patients with proximal colonic adenomas and tumors. Serrated tumors have specific molecular attributes as they do not emanate from initiating mutations in the *APC* tumor suppressor and therefore do not present hyperactivation of the Wnt/β-catenin pathway ^44,45,61,62^. While signature of Lgr5^+^ crypt-base columnar stem cells is enriched in tumors following the conventional adenoma-to-carcinoma sequence ^63,64^, serrated tumors display increased expression of fetal markers ^44^.

Our Gene Set Enrichment Analysis (GSEA) provides a mechanistic bridge between human cohort analyses and studies in organoid models. RNA-seq in organoids revealed a robust and reproducible enrichment of signatures associated with serrated colorectal cancer while signatures of the conventional adenoma-to-carcinoma pathway were not enriched. Conversely, Wnt/β-catenin–related gene sets were negatively enriched, consistent with our observation of reduced expression of canonical Wnt target genes (*e.g.*, *Axin2*, *Lef1*). Importantly, colonic mucosal transcriptomes from CRC patients carrying *cnf1*⁺ bacteria also displayed enrichment of serrated signatures and downregulation of Wnt-associated programs, mirroring the transcriptional changes induced in organoids. These convergent findings strongly support the notion that CNF1-expressing *E. coli* drive a serrated-like trajectory of colorectal tumorigenesis.

Mechanistically, CNF1 activates a RhoA/Rock axis that reprograms Lgr5⁺ adult stem cells into fetal-like progenitors with enhanced stemness potential. This represents, to our knowledge, the first example of a bacterial toxin directly driving epithelial-intrinsic stem cell reprogramming toward a fetal-like state. Involvement of Yap in fetal reprograming has been described in situations of DSS or radiation-induced damages to the adult Lgr5^+^ intestinal stem cell (ISC) population and Diphtheria-toxin mediated conditional ablation of intestinal stem cells ^51,65,66^. Here, we show that Yap/Taz induction downstream of RhoA/Rock mediates the induction of fetal markers. RhoA has been proposed to induce Yap translocation into the nucleus through Rock-mediated actomyosin contractility, impacting the nuclear pore aperture ^67^.

RhoA activation has been implicated in a wide range of cancers, where it regulates key oncogenic processes such as epithelial-to-mesenchymal transition, invasion, and metastasis. In the context of cancer initiation, literature highlights the context-dependent contribution of RhoA signaling to tumorigenesis. Activation of the RhoA/Rock signaling pathway in mouse skin has been shown to promote nuclear accumulation of β-catenin and enhance its transcriptional activity, ultimately leading to epithelial hyperproliferation and tumor development ^68^. Paradoxically, in the intestinal epithelium, chronic β-catenin activation and increased tumor incidence were observed upon RhoA inactivation, through expression of a dominant-negative RhoA mutant ^69^. In gut organoids, we show that CNF1-mediated RhoA activation is associated with decreased Wnt/β-catenin activation, which is consistent with the later publication, and to induction of fetal-like stem cells, a feature of serrated tumorigenesis of the colon.

The organoid model allowed us to unravel RhoA involvement downstream of the CNF1 bacterial toxin in gut epithelial stem cell fate commitment. The transition to fetal-like state could be instrumental to generate cells that are less lineage-restricted to help them evade differentiation cues and allow tumor emergence. Moreover, reactivation of a fetal-like stem cell program brings plasticity to gain growth and survival advantage in specific environments ^70^. In link with these features, the temporal link between developmental reprogramming-mediated plasticity and colorectal cancer initiation was demonstrated in serrated tumors ^44,71^.

In conclusion, we propose that CNF1 toxin-producing *E. coli* contribute to the serrated pathway of colorectal tumorigenesis by reactivating fetal-like stem cell programs in adult Lgr5^+^ stem cells. Moreover, mechanisms we describe at the epithelium level should help to understand versatility of CNF1 action in different animal models of CRC, where stromal and immune contexts differ. The specific enrichment of *cnf1⁺*bacteria in early proximal lesions positions CNF1 as a potential microbial biomarker and therapeutic target in CRC, underscoring the broader role of microbiota-derived virulence factors in shaping distinct molecular subtypes of cancer.

## MATERIALS AND METHODS

### Recruitment of Participants and Collection of Samples

Patients were enrolled with informed consent between 2004 and 2018 in several prospective cohorts named CCR1, CCR2, Vatnimad, DETECT by the endoscopy department at Henri Mondor hospital (APHP, Créteil, France), where patients had been referred for colonoscopy, see for detailed description ^34^ and ClinicalTrials.gov: NCT01270360. The study was approved by the ethical committee of Val de Marne Paris-EST area (no. 10-006 in 2010).

Exclusion criteria for these cohorts were: a history of colorectal surgery due to CRC, familial adenomatous polyposis, Lynch syndrome, infection, inflammatory bowel disease and exposure to antibiotics during the 3 weeks preceding colonoscopy. Carcinoma were classified according to Tumor Node Metastasis (TNM) staging. Clinical parameters of the patients, such as body mass index (BMI), age, sex, and disease history were referenced (Table 2).

Fresh stool samples were collected 2 weeks to 3 days before colonoscopy, prior to bowel cleanse, frozen at −20°C within 4 h and deposited at the Henri Mondor Hospital biobank CRB (Biological Resources Center)^34^; here referred to as cohort FR2. Paired samples of colorectal tumor tissue and homologous normal mucosa (more than 15 cm from the margin of tumor resection) were collected within 30 min after surgical resection and immediately frozen and stored at −80°C until use ^41^; material here referred to as cohort FR3.

### Fecal and tissue DNA extraction

DNA was extracted from the stools using the Wizard Genomic DNA Purification kit (Promega, A1120), modified according to ^72^.

DNA was extracted from tumor tissue and homologous normal mucosa starting from eight 50 μm cryosections of nitrogen frozen tissue using the QIAamp PowerFecal DNA Kit® (Qiagen, 12830-50) following the manufacturer’s instructions with the following modification: 0.1 mm diameter silica beads were added to the lysis solution provided and shaking was performed at maximum speed for 10 minutes in a vibratory shaker.

DNA concentration was measured using the Qubit dsDNA Broad Range assay kit (Invitrogen Q32853).

### Derivation of a CNF1/2 consensus sequence and design of the *cnf1* primers

From previous work in the lab, we identified 6,596 *cnf*-like toxin positive genomes out of 141,243 *E. coli* genomes collected from Enterobase ^18^. From these genomes, were extracted 5,656 complete *cnf*-like toxin encoding genes sequences (3,045 bp). This corresponds to 5,187 sequences of *cnf1* and 469 sequences of *cnf2*. To this, was also added the sequence of *cnf1* from *E. coli* UTI89. Using MAFFT ^73^, we aligned the total of 5,657 *cnf*1/2 nucleic sequences. Next, with Emboss cons we calculated a consensus sequence by setting at 90 % the minimal identity at a site (Supplementary Information 3). Then, primers specific of *cnf*1/2 were designed in the consensus sequence using the NCBI primer tool (Table 4). Positive hits were all sequence verified, they corresponded to *cnf1*.

**Table 4.**
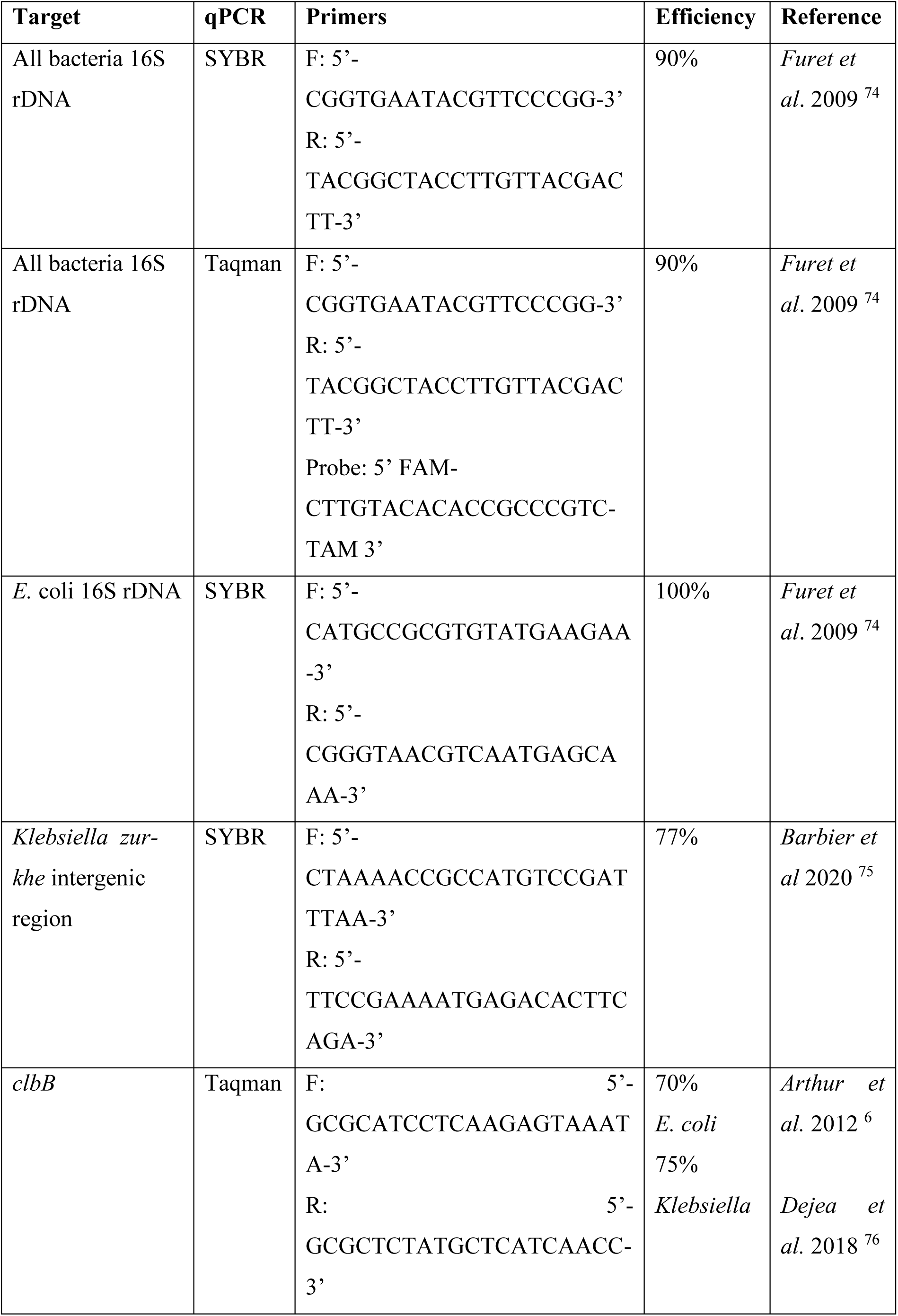
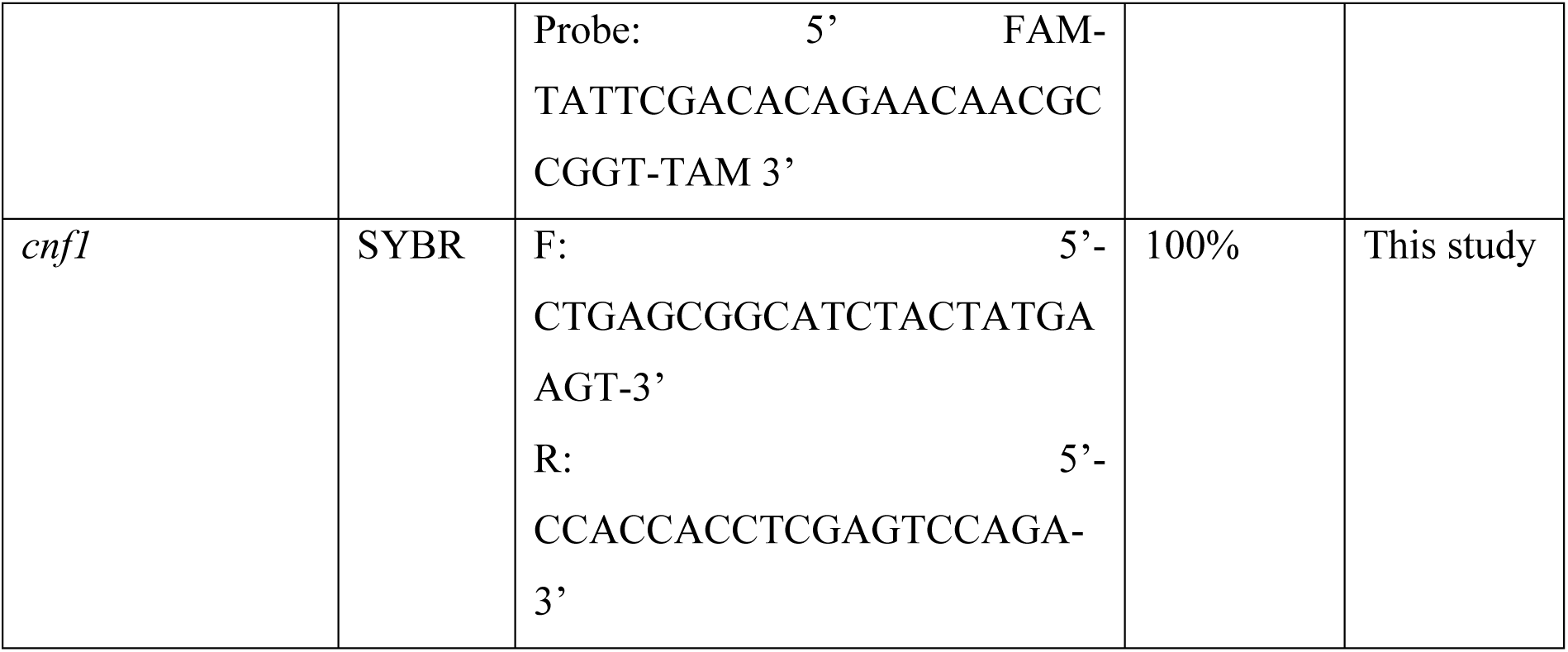
Sequence of the primers, and probes for Taqman detection, used for qPCR detection. Efficiency of the primers was calculated using a 5-fold serial dilution of DNA from *Escherichia coli* UTI89 or *Klebsiella pneumoniae* CIP 82.91T, ATCC 13883.

### qPCR on human fecal and tissue DNA

qPCR reactions were performed on a QuantStudio^TM^ 7 Flex Real-Time PCR System (Applied Biosystems, 4485701). Each sample was run in triplicate. Primers and probes listed in Table 4 were ordered from Sigma. Primers against the colibactin gene *clbB* were used to detect *pks*. Efficiency of the primers was determined using a 5-fold serial dilution of DNA from *Escherichia coli* UTI89 or *Klebsiella pneumoniae* (strain CIP 82.91T, ATCC 13883, obtained from the Collection of Institut Pasteur).

The Master Mix SYBRGreen (Applied Biosystems, 4367659) or the Master Mix Taqman (Applied Biosystems 4440038) were used according to the manufacturer’s instructions. The amount of target gene was determined using the QuantStudio^TM^ Real-Time PCR software (version 1.7.2, Applied Biosystems) and the 2^-ΔCt^ method, with all bacteria 16S rDNA as a calibrator. Some patients were randomly selected, amplicons were purified with the NucleoSpin Gel and PCR Clean-up (Macherey-Nagel™, 740609.50), and the specificity of amplification was validated by Sanger sequencing (Eurofins Cologne).

### WGS data collection

Seven previously published ^40^ colorectal cancer projects with WGS available on the ENA website ^77^ were used for our in-silico analysis (Table 1). We collected metagenomic datasets for each project. This corresponds to sequencing runs from 884 samples in total (382 controls, 116 adenomas and 386 CRC). Associated metadata for each sample was retrieved both from ENA website and from GMrepro ^78^ using the project id as query. Archive generated FastQ files were converted in Fasta file containing nucleic acid information, then reads were mapped with minimap2 ^79^ to the human genome (GRCh38/hg38) and processed with samtools ^80^ for the removal of human sequences contaminations. We used Emboss transeq to translate filtered reads into peptide sequences in the three forward and three reverse frames before screening for *cnf1* and *pks*-island.

### Diamond in house databases creation for CNF1 and *pks*-island protein sequences

We created a collection of CNF-like sequences by mining *E. coli* genomes from Enterobase and UNIPROT database using an HMM profile previously described ^18^, which was designed to capture the entire diversity of *cnf*-like toxin protein family. We next extracted the CNF1-domain of these CNF-like sequences and removed duplicates. We then obtained 122 unique representative CNF1-domain from CNF-like sequences that were used for the construction of the CNF-like toxins database.

*pks*-island from 60 *E. coli* complete genomes were extracted based on the successive alignment and localization with mummer3 ^81^ using UTI89 *pks*-island as the reference sequence. Aligned regions from *pks*-island positive genomes were submitted to Prokka ^82^ for the identification and annotation of coding sequences based on the annotated *pks*-island of *E. coli* UTI89 (ClbA-ClbS) obtained via Islandviewer4 ^83^. Translated coding sequences were then collected and used for the creation of *pks*-island sequences Diamond database.

### Identification of *cnf1* and *pks*-island positive samples

We used Diamond ^84^ to align filtered reads against our reference databases to search for *cnf1* or *pks*-island sequences positive reads. We set the minimal percentage of sequence identity and query coverage to 95% to avoid false positive reads. Only one single hit was retained for each matching read. Each sample with at least one c*nf1*-positive read was considered as *cnf1*-positive sample and samples with at least 10/19 unique *pks*-island sequences were considered as *pks*-island positive.

### Mice

All animal experiments were approved by the committee on animal experimentation of the Institut Pasteur and the National Research Council (APAFIS#26133-202006221228936). C57BL/6 female mice aged 8-17 weeks were used. Mice were either bred at the Institut Pasteur specific pathogen-free animal facility or ordered from Charles River. Lgr5-EGFP-IRES-Cre^ERT2^ (JAXstock #008875) were ordered from The Jackson Laboratories, and crossed with ROSA^mT/mG^Cre reporter mice ^85^ provided by Institut Pasteur to obtain GFP-LGR5;mTmG mice.

### Primary murine intestine organoid culture

Small intestine or colon organoids were cultured as previously described ^33^, independent isolations from mice were performed for each experiment. The duodenum or colon was digested in 10 mM EDTA (Sigma, E5134-500G). Intestinal crypts were seeded at 10-15 crypts/μl in a 2:1 Matrigel (Corning, 356231) : small intestine organoid culture medium for duodenal crypts, or in pure Matrigel for colonic crypts. Small intestine organoid culture medium (ENR) was added consisting of advanced DMEM F12 (Gibco, 12634-010) supplemented with Glutamax (Gibco, 35050038), 100 units/ml penicillin-streptomycin (Gibco, 15140122), Hepes (Gibco, 15630056), N2 (Gibco, 17502-048), B27 (Gibco, 12587-010), 50 ng/ml murine EGF (Peprotech, 315-09), 100 ng/ml Noggin (Peprotech, 250-38) and 500 ng/ml mouse R-spondin 1 (R&D, 3474-RS-050). Colon organoid culture medium (ENRW) comported 1 μg/ml R-spondin 1, and was further supplemented with 200 ng/ml Wnt3a (Peprotech, 315-20), 3 μM CHIR99021 (Sigma, 1046), 10% decomplemented fetal bovine serum (Cytiva, SV30160.03), and 10 μM Y-27632 for the first 24h (Sigma, Y0503).

### Organoid passaging

Organoids were collected on ice in organoid harvesting solution (Cultrex, 3700-100-01), disrupted by forceful pipetting, pelleted for 5 min at 100 g at 4°C, washed once in advanced DMEM F12 and resuspended in pure Matrigel (Corning, 356231).

### Organoid stimulation

Small intestine organoids were stimulated 4 days after primary organoid culture. Colon organoids were cleaned and passaged 1-3 times until the culture achieved appropriate growth and morphology, and stimulated 3 or 4 days after the last passaging.

Unless otherwise specified, organoids were stimulated with 0.1 nM CNF1 WT or CNF1 C866S purified as described in ^28^. Where indicated, 3 μM of an inhibitor of Yap-Tead interaction (verteporfin, Sigma, SML0534-5MG), and 10 μM of a Rock inhibitor (Y-27632, Sigma, Y0503-1MG) were added.

### Organoid culture from single cells

Organoids were either untreated or stimulated for 24 h with 0.1 nM CNF1, and dissociated for 5 min at 37°C in TrypLE™ Express (Gibco, 12605010). Organoids were washed once in advanced DMEM F12 (Gibco, 12634-010), filtered with a 40 μm strainer (Falcon, 352340), and 15,000 cells were plated in 30 μl drops of Matrigel (Corning, 356231). Small intestine ENR organoid culture medium was supplemented with 200 ng/ml Wnt3a (Peprotech, 315-20) and 1 μM Jagged-1 (AnaSpec, 188-204).

### RNA extraction

Organoids were collected in cold PBS, washed once, and RNA was extracted using the RNeasy mini kit (Qiagen, 74104) or the RNeasy micro kit (Qiagen, 74004) according to the manufacturer’s instructions. RNA concentration was quantified using a DeNovix DS-11 FX+ spectrophotometer.

### RNA sequencing

For RNA-seq on organoids, Routine DNase treatment was performed on organoid RNA with the TURBO DNA-free™ Kit (Invitrogen, AM1907) according to the manufacturer’s instructions. RNA concentration was quantified using Qubit (Invitrogen, v 4). The RNA Integrity Number was measured following the manufacturer’s instructions, using the Bioanalyzer 2100 with the 2100 Expert software (Agilent, v B.02.12), the RNA nano chips and the RNA 6000 Nano Kit (Agilent, 5067-1511).

Libraries were built using an Illumina Stranded mRNA library Preparation Kit (Illumina) following the manufacturer’s protocol. To decrease the remaining quantity of primer dimers, two purifications with Ampure XP beads (Beckman Coulter, A63882) were necessary. Sequencing was performed on a NextSeq2000 (Illumina) to obtain 117 base single-end reads. RNA-seq count data were analyzed using R software (version 4.4.2) and RStudio (version 2025.05.0+496) with DESeq2 (package version 1.44.0) for differential expression analysis. Functional enrichment analysis was performed using Gene Set Enrichment Analysis (GSEA) ^86^ implemented in package cluster Profiler (version 4.16.0) with a composite ranking score calculated as:

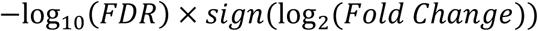

with random jitter (± 10^−6^) added to break ties and averaged scores for genes with multiple transcripts. Eight gene set collections were tested including six from MSigDB v 2024.1 (mouse): Hallmark, Curated gene sets (M2), Regulatory targets from GTRD (M3), GO Biological Processes (M5), Cell type signatures (M8), and KEGG pathways, plus a custom collection specific to this study (“curated genesets”, Supplementary Information 2). GSEA was performed using an adaptive multilevel splitting Monte Carlo approach (ε = 0). Benjamini-Hochberg correction was applied for multiple testing in both differential expression and functional enrichment analyses. Gene sets were considered significantly enriched when the normalized enrichment score (NES) exceeded an absolute value of 1.0 and the false discovery rate (FDR) was < 0.05.

RNAseq on FR3 cohort patients is described in ^41^, with the following modification: normalized counts were generated using the voom procedure ^87^ implemented in the limma package (v 3.52.4) ^88^.The GSEA was performed using fgsea R package ^89^ based on the log(fold change) and on curated gene sets (Supplementary Information 2).

### RT-qPCR

Total RNA was reverse-transcribed into cDNA with the Super script reverse transcriptase SS IV RT (Invitrogen, 18090050) using a BiometraTOne 96 G (Analytik Jena, 846-2-070-301). Power SYBRGreen PCR master mix (Applied Biosciences, 4367659) was used to amplify cDNA according to the manufacturer’s instructions. Abundance was measured using QuantStudio 7 real-time PCR system (Applied Biosystems, 4485701) and analyzed on QuantStudio™ Real-Time PCR Software (Applied Biosystems, v1.7.2). cDNA abundance was calculated using the 2^-ΔΔCt^ method, with *Gapdh* as a normalization control.

All primer pairs listed in Table 5 were validated by amplicon sequencing (Eurofins, Cologne) after purification using the NucleoSpin Gel and PCR Clean-up Mini kit (Macherey Nagel, 740609.50) following the manufacturer’s instructions.

**Table 5.**
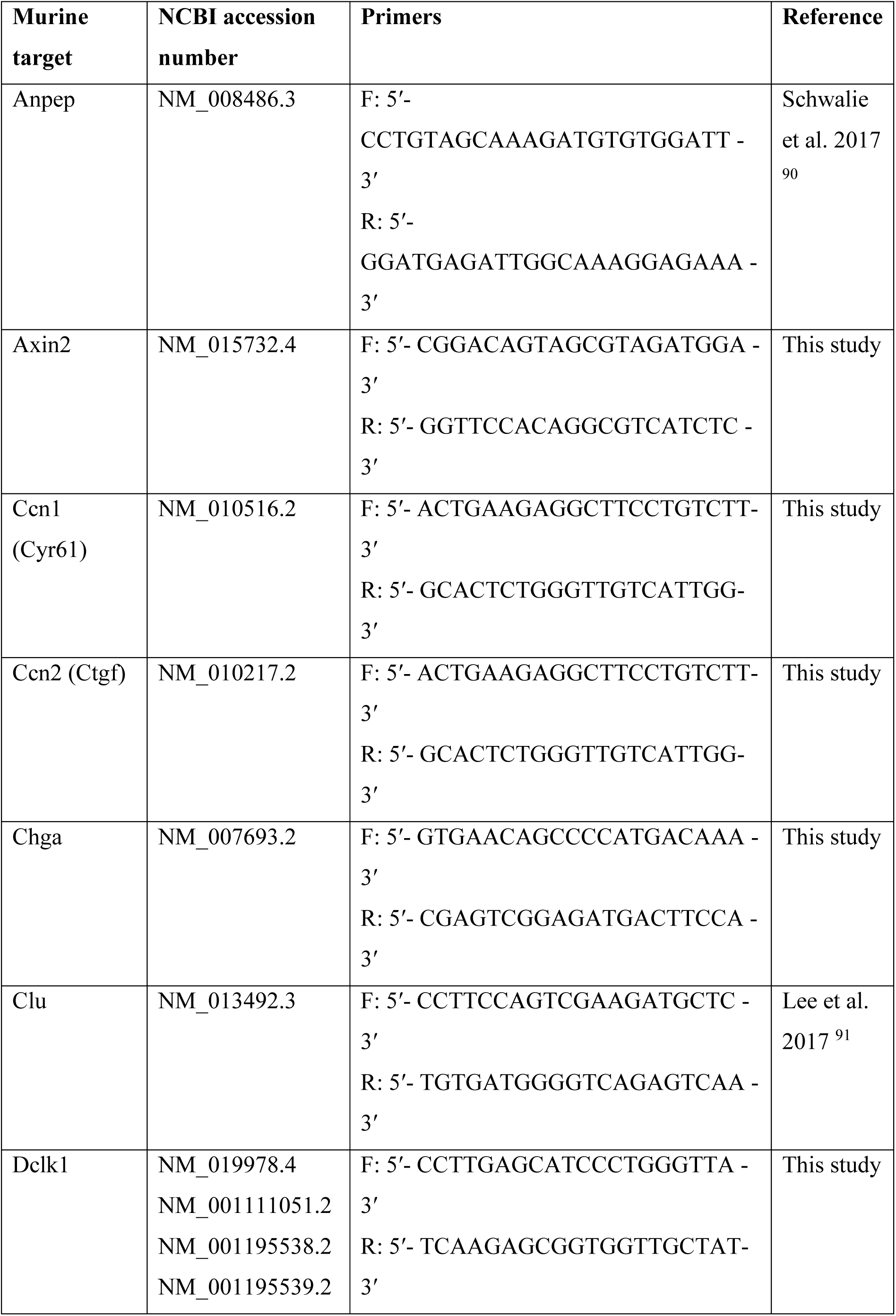
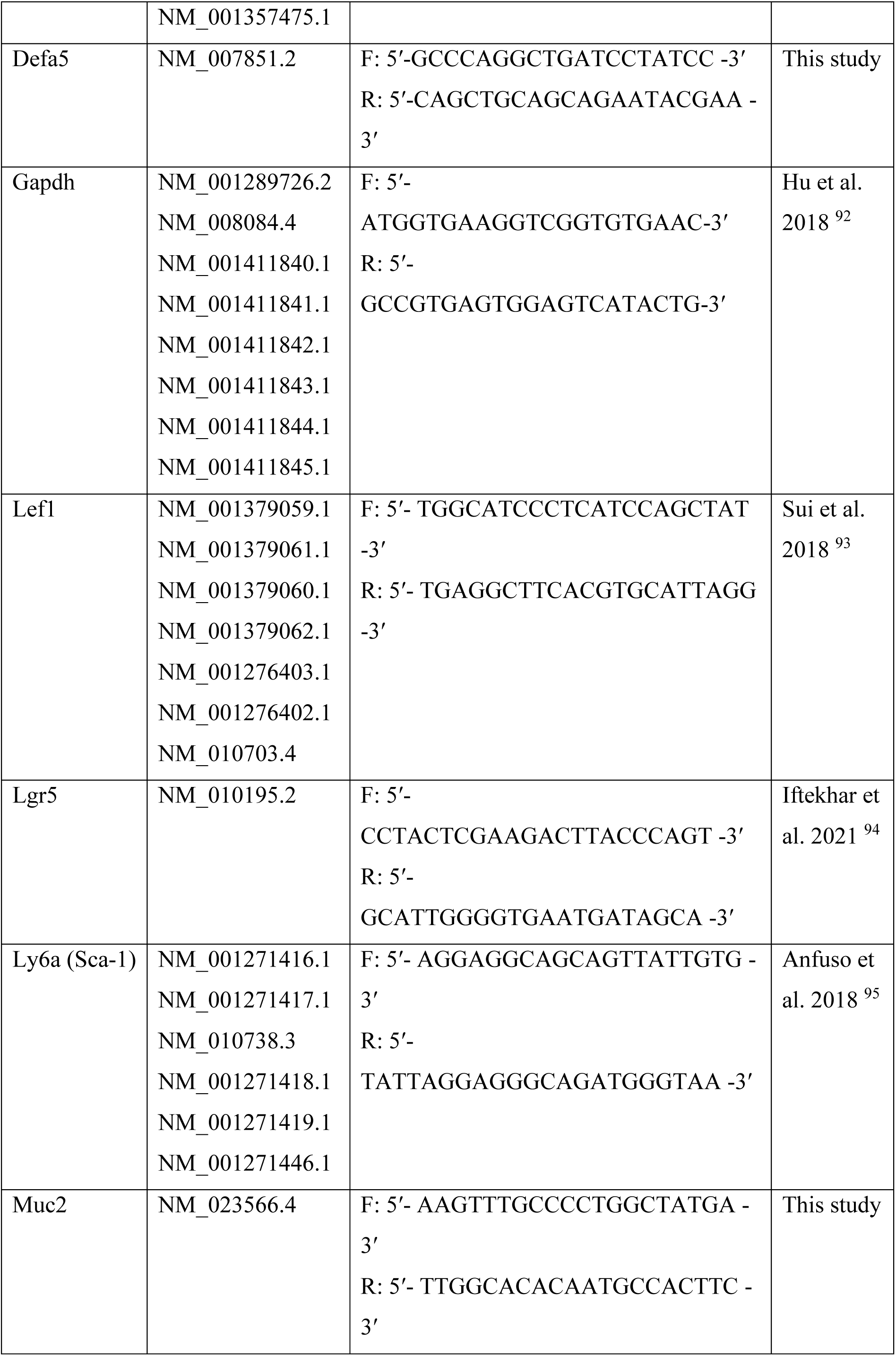
List of the primers used for RT-qPCR detection on murine material.

### Flow cytometry

Lgr5-GFP organoids were dissociated in TrypLE™ Express (Gibco, 12605010) and stained with anti-Sca-1 antibody (E13-161.7 clone, Biolegend, 122501) and 0.1 μg/ml DAPI (Sigma, D9542). Sorting was performed with the MoFlo Astrios (Beckman Coulter, France) at 25 PSI with a 100 nm nozzle at approximately 2000 events per second. GFP fluorescence was read with the 488 laser (576/21 band pass) and Sca-1 with the 561 laser (614/20 band pass), and analyzed with FlowJo (BD, v 10.10.0). Cell doublets and dead cells were excluded from the analysis.

### Organoid immunolabeling and imaging

Organoids were fixed in 4% PFA (Electron Microscopy Sciences, 15710) at 4°C overnight. For imaging, permeabilization was performed either in 1% SDS for 30 min at 37°C followed by extensive washing and 15 min at RT in 0.5% Triton X-100 (Sigma, T9284) for Yap translocation assay, or in 0.5% Triton X-100 for 1 h at RT for Ki67 and Sca-1 staining. Organoids were blocked in 3% BSA for 1 h at RT, and incubated overnight at 4°C in anti-Yap (Santa Cruz, sc-101199), anti-Ki67 (eBiosciences, 50-5698-82), anti-Sca-1 (Biolegend, 122501), or anti-E-cadherin (CST, 14472) primary antibody diluted in 0.2% Triton X-100 1% BSA, incubated for 1 h at RT in secondary antibody, and incubated 10 min in 1 µg/ml DAPI (Sigma, D9542). After PBS washes, organoids were mounted in Permafluor (Epredia, TA-030-FM). Images were acquired either with a Confocal Opterra microscope with a Zeiss Axio Observer using a 63x/1.40 oil M27 objective and the Prairie View software (version 5.3), or with a Nikon Ti2E microscope equipped with a Yokogawa CSU-W1 spinning disk using a 63x 1.4 OIL WD 0.13 objective, with a Photometrics Prime 95B camera and the NIS Element software (version 5.42.06). For Ki67 and Sca-1 staining, images were analyzed with Fiji (ImageJ2-win64). Organoid cells with Ki67 nuclear mean fluorescence intensity (MFI) above 2000 were considered as positive.

### Quantification of Yap nuclear translocation

Images were analyzed using Cell Profiler (v 4.2.6). Background illumination was corrected for each channel using a block size of 20 and Gaussian smoothing. Nuclei were identified as primary objects on Gaussian-filtered DNA images, with an expected diameter of 40–70 pixels; objects outside this range or touching the image borders were excluded. Thresholding employed a global approach using the Minimum Cross-Entropy method, with a smoothing scale of 1.35 and bounds of 0.005–1.0. Clumped nuclei were separated based on intensity, with smoothing parameters and minimum distances between local maxima determined automatically. After thresholding and declumping, nuclei masks were filled to remove holes. To avoid capturing cytosolic signal, nuclei masks were shrunk by 5 pixels, and cytosolic masks were defined as a 2-pixel expansion around the nuclei. The percentage of Yap signal within nuclei was calculated as the median Yap intensity in the shrunken nuclei divided by the sum of median intensities in the shrunken nuclei and cytosol.

### RNAscope

Organoids were grown in an 18-well µ-Slide (ibidi, 81817) and fixed overnight at 4°C in 4% PFA. All washes were performed in PBS 0.2% Tween 20 µg/ml Heparin. Organoids were delipidated in 4% SDS 200 mM Boric Acid pH7 for 2 h at 40°C, treated six times with PBS 0.5% Triton for 5 min at RT, washed twice, and incubated in PBS 0.2%Tween 3%H2O2 for 10 min at RT. After 2 washes, RNAscope hybridization was performed with the Lgr5 C2 probe and 650 TSA Vivid Dye according to the manufacturer’s instructions (ACD, 323110). Samples were visualized with a STELLARIS 8 inverted microscope (Leica Microsystems, Mannheim, Germany). A super continuum white light laser tunable between 440 and 790 nm and a 405 laser were used for excitation and focused through an HC PL APO CS2 20x NA 0.75 dry objective. Emission signals were captured with Power HyD Detectors. The system was controlled with Leica Application Suite (LAS) X v4.8.1.29271 software. The excitation wavelength and detection window are given in parentheses for each fluorophore: DAPI (405 nm; 427–475 nm); AlexaFluor488 (499 nm; 504–564 nm); AlexaFluor594 (590 nm; 595–650 nm); TSA Vivid Dye (654 nm; 659–750 nm).

### Statistical analysis

Statistical analysis was performed using R (4.2.2) and RStudio (2024.12.0+467). To perform pairwise comparison of the proportion of *cnf1*, *pks*-island, and *Klebsiella* positive samples between the three groups (control, adenoma and CRC), we either used the Chi-2 test for samples with n > 5 or the Fisher test (function pairwise.prop.test from the R stats package (v 4.3.0) or function pairwise.fisher.test from the R fmsb package (v 0.7.6), respectively). To test the significant difference of relative abundance of *E. coli* in samples from the different groups we performed non-parametric Wilcoxon rank-sum test with Benjamini-Hochberg correction. Statistical analysis of RTqPCR comparisons and percentages determined by FACS or immunofluorescence staining was performed by generalized linear regression using independent experiments as random effects with Tukey’s correction for multiple testing.

## ACKNOWLEDGEMENTS

This work was financially supported by institutional fundings from INSERM and CNRS, Institut Pasteur, Université Paris-Est Créteil-PHRC 2011 VATNIMAD, from French National Research Agency-ANR (ExPECtation, ANR-21-CE15-0006), Fondation ARC (PJA 20191209650), Ligue contre le cancer (RS20/75-63), Institut Pasteur Cancer Initiative program, ITMO Cancer AVIESAN (National Alliance for Life Sciences & Health) within the framework of the Cancer Plan [HTE201601] supported by INCA, and executed within the frame of the Oncomix research program between Institut Pasteur and AP-HP. CDSN ENS Lyon and FRM (FDT202404018669) doctoral fellowships to DL, Pasteur-Roux-Cantarini post-doctoral fellowship to EB.

We are grateful to Pr P. Sansonetti for his invaluable guidance and for initiating the Oncomix framework, a pioneering contribution that was instrumental to this work.

We warmly thank M.A. Nahori for technical assistance with animals, and our short-term interns L Lamy and E. Glading for their valuable assistance during this project. We acknowledge Bruno Périchon and Shaynoor Dramsi for sharing information on CCR patients analysis, and we warmly thank all patients that participated in this study. We acknowledge URC-est, St Antoine Hospital and CRB, Henri Mondor Hospital APHP for their involvement in clinical trials.

We thank the Institut Pasteur facilities for their support and expertise : UTechS Photonic BioImaging (Imagopole), C2RT, supported by France BioImaging, ANR-24-INBS-0005-FBI-BIOGEN; Investments for the Future Région Ile-de-France (DIM1HEALTH) and the French Government Investissement d’Avenir Programme Laboratoire d’Excellence ‘Integrative Biology of Emerging Infectious Diseases’ (ANR-10-LABX-62-IBEID), and acknowledge support from Christelle Travaillé for the use of the Nikon Ti2E spinning disk microscope, and Julien Fernandes for the use of the Stellaris 8 microscope. Biomics Platform, C2RT, supported by France Génomique (ANR-10-INBS-09) and IBISA and acknowledge support from Marc Monot, Elodie Turc, Laure Lemée, Etienne Kornobis and Rania Ouazahrou. We thank for their valuable support Thomas Obadia from Biostatistics hub, Vincent Guillemot and Nathalie Lehman from Bioinformatics Hub, Pierre Henri Commere from the Flow cytometry facility.

## LEGENDS TO SUPPLEMENTARY FIGURES

**Figure S1.**
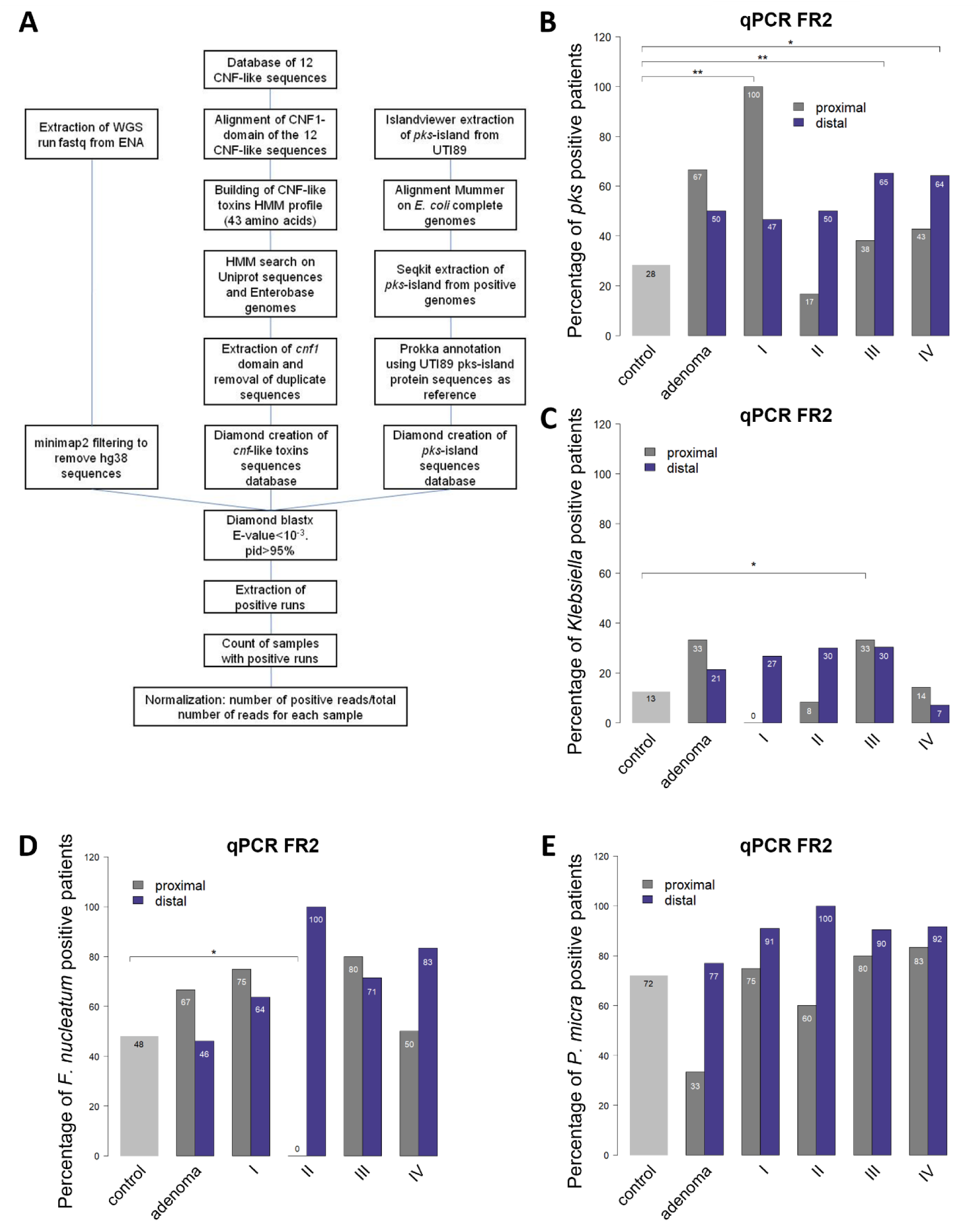
Increased prevalence of CNF1 and Colibactin toxins-encoding genes in colorectal cancer-associated microbiota. (A) Pipeline of custom database creation for *cnf1* and *pks* reference sequences and Diamond analyses. (B) Percentage of patients from the FR2 cohort with fecal carriage of *pks* depending on the stage and tumor location. Chi2 and Fisher’s exact test were used to determine statistical significance. ^∗∗^ *p* < 0.01, ^∗^ *p* < 0.05. (C) Percentage of patients from the FR2 cohort with fecal carriage of *Klebsiella* species. Fisher’s exact test was used to determine statistical significance. ^∗^ *p* < 0.05. (D, E) Percentage of patients from the FR2 cohort with fecal carriage of *Fusobacterium nucleatum* (F) and *Parvimonas micra* (G) depending on the stage and tumor location. Fisher’s exact test was used to determine statistical significance. ^∗^ *p* < 0.05.

**Figure S2.**
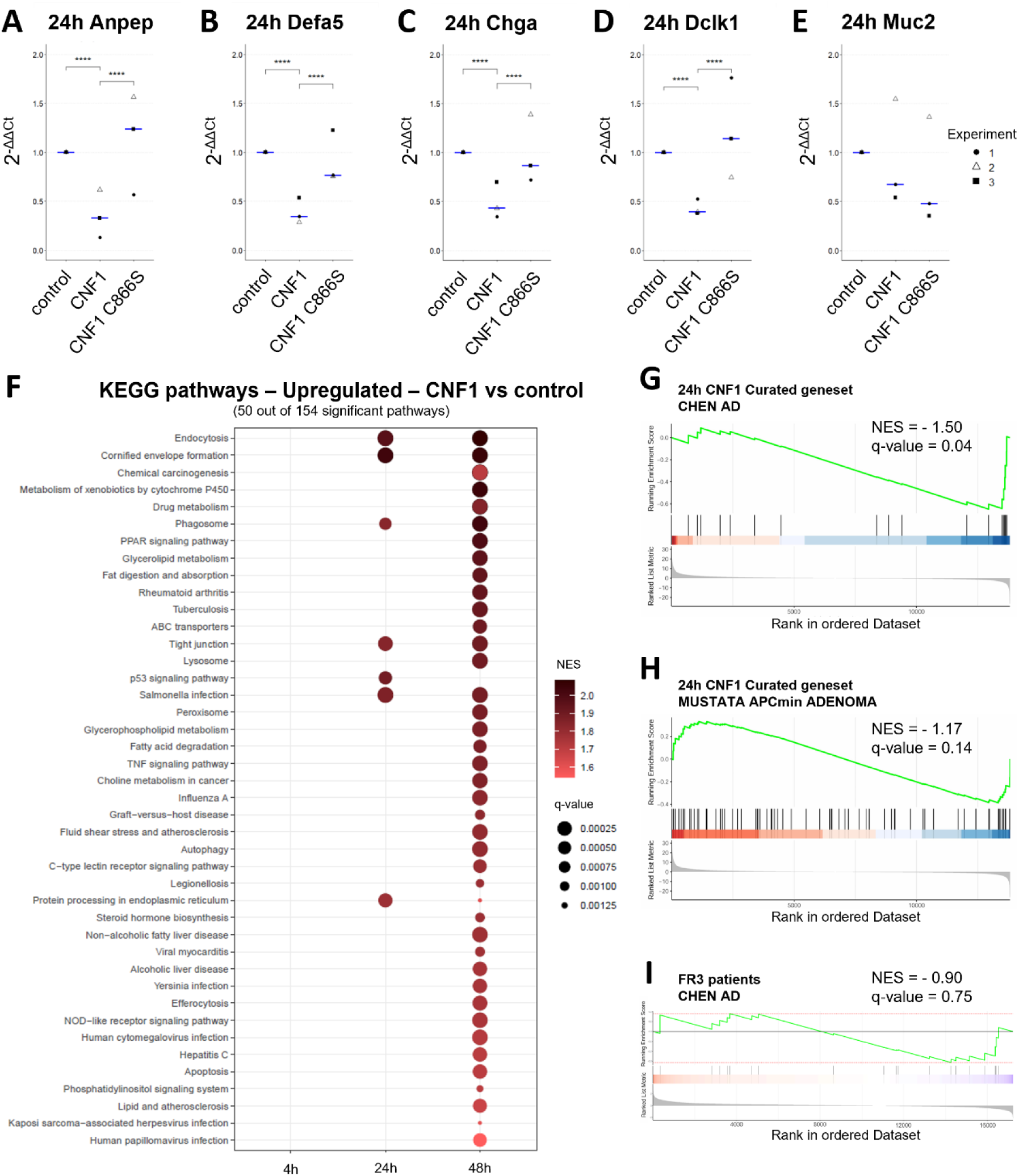
The CNF1 toxin of *E. coli* triggers a serrated pathway transcriptional signature of carcinogenesis in intestinal organoids. (A-E) RTqPCR showing expression of markers of enterocytes (*Anpep*) (A), Paneth cells (*Defa5*) (B), enteroendocrine cells (*Chga*) (C), Tuft cells (*Dclk1*) (D), and goblet cells (*Muc2*) (E) after 24h of stimulation with 0.1 nM of CNF1 (WT or C866S), with median in blue (n=3 independent experiments). Statistical significance was determined by generalized linear regression using independent experiments as random effects, with Tukey’s correction. ^∗∗∗∗^ *p* < 0.0001. (F) KEGG pathways most enriched in CNF1-treated organoids according to normalized enrichment score and q-value by DESeq analysis with Benjamini-Hochberg correction (n=3 independent experiments). (G-H) GSEA analysis in organoids after 24h of 0.1 nM CNF1 stimulation showing the Chen signature of conventional colon adenoma (G) and the Mustata signature of *Apc^Min/+^* adenomas (H). Statistical significance was determined by Monte Carlo analysis with Benjamini-Hochberg correction (n=3 independent experiments). (I) GSEA analysis of samples from the FR3 cohort with *cnf1* tissue carriage compared to samples without *cnf1* carriage, showing the Chen signature of conventional colon adenoma. Statistical significance was determined by Monte Carlo analysis with Benjamini-Hochberg correction. Not significant

**Figure S3.**
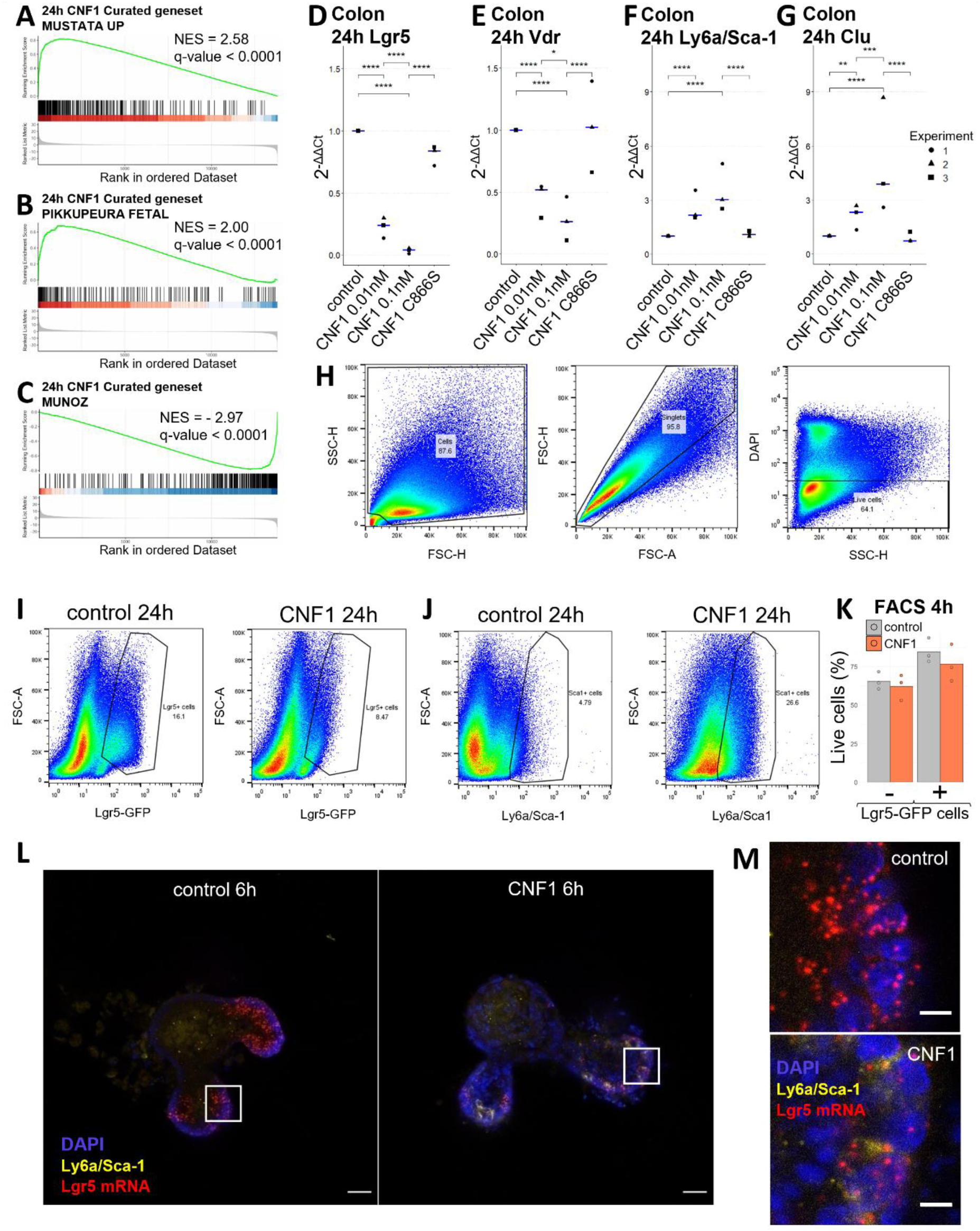
The CNF1 toxin of *E. coli* activates a fetal-like transcriptional program in stem cells. (A-C) GSEA analysis of the Mustata signature of fetal intestinal self-renewing spheroids (A), the Pikkupeura signature of fetal proximal mouse small intestine (B), and the Muñoz signature of Lgr5^+^ cells (C) after 24h of 0.1 nM CNF1 stimulation. Statistical significance was determined by Monte Carlo analysis with Benjamini-Hochberg correction (n=3 independent experiments). (D-G) RTqPCR showing expression of the Lgr5^+^ stem cell markers *Lgr5* (C) and *Vdr* (D), and the fetal markers *Ly6a*/Sca-1 (E) and *Clu* (F) in colonic organoids after 24h of CNF1 stimulation, with median in blue (n=3 independent experiments). Statistical significance was determined by generalized linear regression using independent experiments as random effects, with Tukey’s correction. ^∗∗∗∗^ *p* < 0.0001, ^∗∗∗^ *p* < 0.001, ^∗∗^ *p* < 0.01, ^∗^ *p* < 0.05. (H) Gating strategy of flow cytometry analyses. Cells were first gated based on forward scatter (FSC) and side scatter (SSC) parameters to exclude debris. Singlets were then identified using FSC-Area versus FSC-Height to eliminate cell aggregates. Live cells were selected by excluding dead cells stained with DAPI. (I-J) Representative flow cytometry profile of Lgr5-GFP (E) and Ly6a/Sca-1 (F) expression in live cells sorted from small intestinal organoids stimulated 24h with 0.1 nM of CNF1. (K) Percentage of live cells sorted by flow cytometry after 4h of 0.1 nM CNF1 stimulation. Data are represented as a mean (n=3 independent experiments). Statistical significance was determined by generalized linear regression using independent experiments as random effects, with Tukey’s correction. (L-M) Immunofluorescence staining of Ly6a/Sca-1 in small intestinal organoids after 6h of 0.1 nM CNF1 stimulation. Nuclei are stained with DAPI and Lgr5 mRNA with a RNAscope probe. Scale bar: (L) 20 µm or (M) 5 µm.

**Figure S4.**
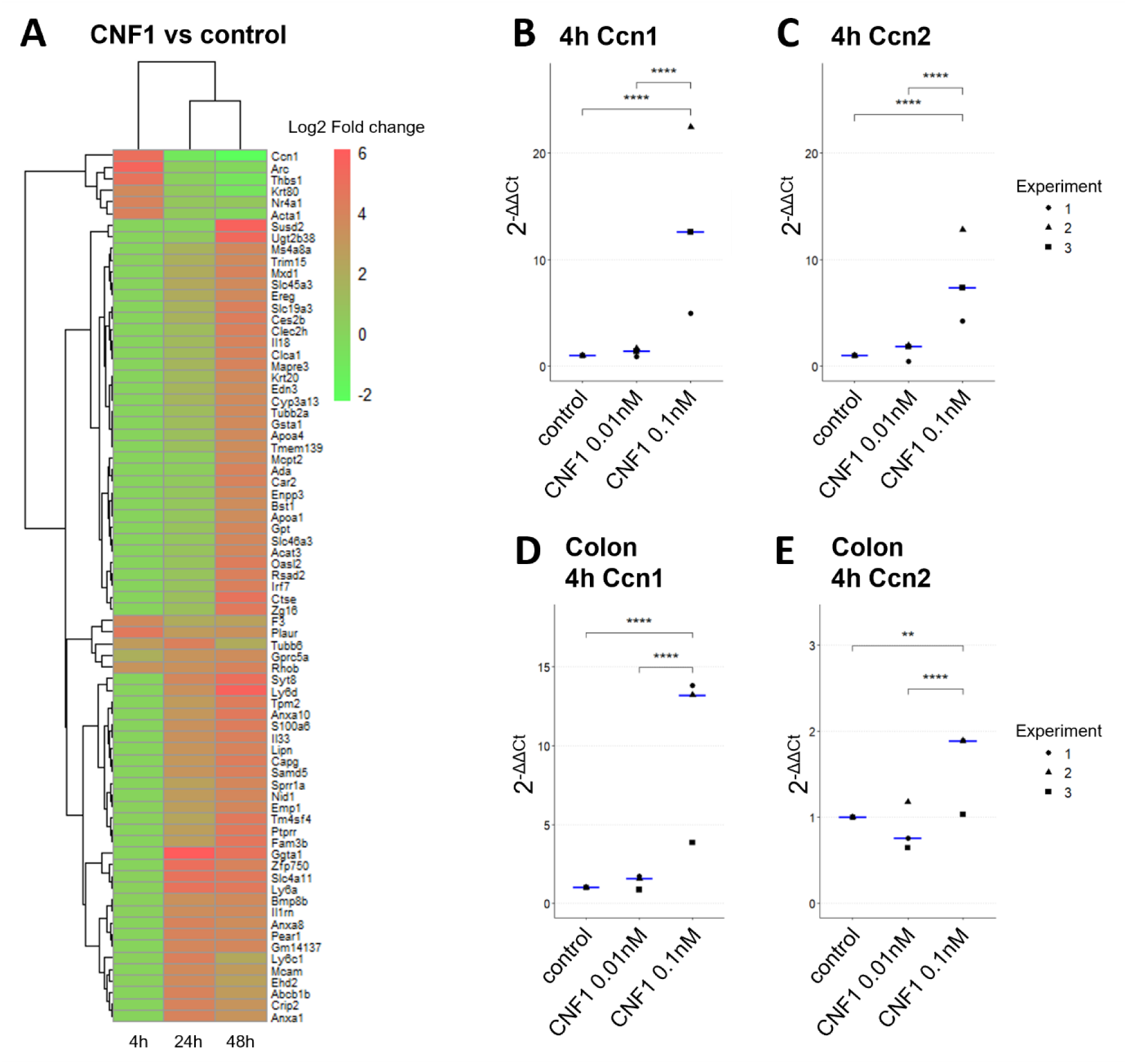
Yap/Taz activity is enhanced by CNF1 in intestinal organoids, in a RhoA/Rock-dependent mechanism. (A) Heatmap showing the log2FC of genes from previously described effectors of the Yap pathway and the Gregorieff Yap signature. Organoids treated with 0.1 nM of CNF1 are compared to non-treated controls. Differential analysis and statistical tests were performed with DESeq, with Benjamini-Hochberg correction (n=3 independent experiments). Genes with q-value < 0.0001 and |log2FC| > 3.5 at 4h, 24h, or 48h were displayed. (B-C) RTqPCR showing expression of the Yap effectors *Ccn1*/*Cyr61* (B) and *Ccn2*/*Ctgf* (C) in small intestinal organoids after 4h of CNF1 stimulation, with median in blue (n=3 independent experiments). Statistical significance was determined by generalized linear regression using independent experiments as random effects, with Tukey’s correction. ^∗∗∗∗^ p < 0.0001. (D-E) RTqPCR showing expression of *Ccn1*/*Cyr61* (D) and *Ccn2*/*Ctgf* (E) in colonic organoids after 4h of CNF1 stimulation, with median in blue (n=3 independent experiments). Statistical significance was determined by generalized linear regression using independent experiments as random effects, with Tukey’s correction. ^∗∗∗∗^ p < 0.0001, ^∗∗^ p < 0.01.

